# Mutagenic ligation of polβ mismatch insertion products during 8-oxoG bypass by LIG1 and LIG3α at the downstream steps of base excision repair pathway

**DOI:** 10.1101/2024.10.23.619805

**Authors:** Kar Men Lee, Erick Castro, Jacob E. Ratcliffe, Melike Çağlayan

## Abstract

Base excision repair (BER) maintains genome integrity by fixing oxidized bases that could be formed when reactive oxygen species attack directly on the DNA. We previously reported the importance of a proper coordination at the downstream steps involving gap filling by DNA polymerase (pol) β and subsequent nick sealing by DNA ligase (LIG) 1 or 3α. Yet, how the fidelity of 8-oxoG bypass by polβ affects the efficiency of ligation remains unclear. Here, we show that LIG1 can seal nick products of polβ after both dATP and dCTP insertions during 8- oxoG bypass, while ribonucleotide insertions completely diminish the repair coordination with both ligases, highlighting a critical role for nucleotide selectivity in maintaining BER accuracy. Furthermore, our results demonstrate that LIG3α exhibits an inability to ligate nicks of polβ dCTP:8-oxoG insertion or with preinserted 3’-dC:8-oxoG. Finally, AP-Endonuclease 1 (APE1) proofreads nick repair intermediates containing 3’-dA/rA and 3’-dC/rC mismatches templating 8-oxoG. Overall, our findings provide a mechanistic insight into how the dual coding potential of the oxidative lesion and identity of BER ligase govern mutagenic *versus* error-free repair outcomes at the final steps and how the ribonucleotide challenge compromises the BER coordination leading to the formation of deleterious repair intermediates.

## Introduction

Oxidative stress caused by reactive oxygen species (ROS) such as superoxide and hydroxyl radicals which arise during metabolic processes in cells and by the effect of environmental exposures such as ionizing radiation is the major threat to genome stability (1). The oxidation of DNA bases by ROS can occur directly in the genomic DNA or indirectly within the nucleotide pool (2). Such oxidative lesions can cause mutations or cell death if they are not efficiently repaired (3). Guanine base is more susceptible to being oxidized by ROS compared to the other four DNA bases due to having the lowest oxidation potential, and guanine oxidation, a phenomenon that underlies age-related disorders including cancer and neurological diseases, leads to 7,8-dihydro-8-oxo-guanine (8-oxoG) formation (4,5).

The coding potential of 8-oxoG is dictated by *anti-* and *syn*-conformations of the oxidized base, which plays a pivotal role in determining its mutagenic potential (6). The equilibrium between these conformations can be influenced by various factors such as cellular environment and presence of specific DNA polymerase (7–10). In the -*syn* conformation, 8-oxoG pairs with adenine through a Hoogsteen base pair leading to a G-T transversion mutation during replication especially in the cancer genome (11). This is caused due to spatial arrangement of the -*syn* conformation of 8-oxoG that closely resembles that of thymine, prompting the polymerase to insert adenine, which constitutes a significant source of oxidative damage-related mutagenesis in cells (12–14). Conversely, 8-oxoG in the -*anti* conformation can pair with cytosine through traditional Watson-Crick base pairing as an unoxidized guanine, forming a non-mutagenic DNA lesion (14–16).

The major defense mechanism against accumulation of 8-oxoG is base excision repair (BER), preventing the mutagenic and lethal consequences of oxidative DNA damage generated by endogenous ROS and environmental toxicants (17–19). The BER pathway involves a series of sequential enzymatic steps and requires a tightly coordinated function of the repair proteins (20,21). The repair pathway begins with a lesion-specific DNA glycosylase that removes a damaged base resulting in an abasic or AP-site in double-stranded DNA (17). This lesion is then recognized by AP endonuclease 1 (APE1) which cleaves the phosphodiester backbone generating a repair intermediate with a single-strand break containing 3’-OH and 5’- deoxyribose phosphate (5’-dRP) groups (18). As a major BER polymerase, DNA polymerase (pol) β removes the 5’-dRP group and then catalyzes a template-based gap filling DNA synthesis by incorporating a single nucleotide into a gap repair intermediate, resulting in the formation of a nick with 3’-OH and 5’-PO4 ends (22). Finally, DNA ligase (LIG) 1 or LIG3α seals the resulting nick repair product by catalyzing a phosphodiester bond formation to join DNA ends to complete the BER pathway (23).

Structural and biochemical studies have extensively characterized the early steps of the BER pathway, including the removal of damaged base by DNA glycosylases (19). 8-oxoguanine DNA glycosylase-1 (OGG1) is the primary enzyme that specifically recognizes and excises 8- oxoG when paired with cytosine by flipping the damaged base out of the DNA helix and cleaving the N-glycosidic bond, creating an AP site that is then further processed by subsequent proteins in the repair pathway (24,25). When 8-oxoG escapes repair, there is a high probability that replication will result in adenine insertion. Thus, the cell also codes for a DNA glycosylase, MutY homolog (MYH or MUTYH), which recognizes and removes adenine paired with 8- oxoG that adopts its mutagenic -*syn* conformation by excising the mispaired adenine (26). Additionally, MutT in bacteria and MutT homolog 1 (MTH1) in human hydrolyze oxidized nucleotides, particularly 8-oxoGTP to its monophosphate form, thereby preventing their incorporation into the genome by DNA polymerases (27). Furthermore, cell studies have identified MTH1 inhibition as an effective tumor-suppressive strategy (28). Together, the coordinated actions of OGG1, MYH, and MTH1 effectively prevent accumulation of oxidative damage from causing mutations and contributes to maintaining the fidelity of the genetic code (29,30).

During gap filling step of BER pathway, polβ can incorporate 8-oxodGTP in both -*syn* and - *anti* conformations opposite adenine and cytosine, respectively (31). We previously reported that the nick repair intermediate with 3’-8oxodGMP inserted by polβ confounds the next ligation step by LIG1 and LIG3α, resulting in ligase failure and formation of abortive ligation products with 5’-adenylate (32,33). When polβ encounters the oxidative damage lesion on the template position, the polymerase exhibits similar efficiency for dATP and dCTP insertions opposite template 8-oxoG, resulting in mutagenic and error-free gap filling, respectively, as shown in structure and kinetics studies of the enzyme (34–39). Particularly, the processing of polβ dATP:8-oxoG insertion products could lead to mutagenic repair that can be a cellular burden during times of elevated metabolic or environmental stress. Due to the higher abundance of ribonucleotide triphosphates (rNTPs) relative to deoxyribonucleotide triphosphates (dNTPs) in human cells, repair and replication polymerases incorporate rNTPs frequently into the genome (40). Most recently, we reported how the ribonucleotide insertion can adversely impact gap filing function of polβ leading to repair pathway discoordination with the BER ligases (41–43). Although these studies emphasize the importance of polβ in modulating the repair of a base lesion, particularly oxidative DNA damage, it remains unknown how polβ 8-oxoG bypass affects accuracy of the repair pathway coordination at the downstream steps where BER ligases seal resulting nick product.

In the present study, to further characterize the molecular determinants of faithful repair pathway coordination at the final steps of the BER pathway, we questioned the impact of polβ mismatch or ribonucleotide incorporation during 8-oxoG bypass on the efficiency of DNA ligation by LIG1 and LIG3α. For this purpose, we investigated the ligation efficiency after polβ dATP or rATP and dCTP or rCTP mismatch insertions opposite template 8-oxoG by both BER ligases. In addition to these coupled assays that monitor polβ and DNA ligase activities simultaneously, using the nick DNA substrates with preinserted 3’-dA/rA:8-oxoG and 3’- dC/rC:8-oxoG, we investigated end joining abilities of LIG1 and LIG3α in the ligation assays to understand how 8-oxoG on a template position in the absence and presence of a single ribonucleotide at 3’-end of nick DNA affects nick sealing efficiency of both BER ligases.

In addition to its main AP-endonuclease role, via proofreading exonuclease activity, APE1 can correct the errors that polβ made during DNA synthesis step of BER pathway (19). In our previous studies, we demonstrated that APE1 can remove a 3’-mismatched or damaged base from the nick repair products and functionally coordinates with both BER ligases during proofreading coupled to nick sealing depending on the base-pairing architecture of the repair intermediates (44–46). In the present study, we further interrogated the role of APE1 during 8- oxoG lesion bypass by polβ and subsequent nick sealing by LIG1 and LIG3α. For this purpose, we investigated 3’-5’ exonuclease activity of APE1 for the removal of 3’-mismatched base (dA or rA and dC or rC) from the nick DNA substrates containing 8-oxoG on template position and monitored coordination between APE1 and BER ligases (LIG1 and LIG3α) during a mismatch removal coupled to ligation in the same reaction.

Overall, our study provides a mechanistic insight into how 8-oxoG bypass by polβ impacts next ligation step by LIG1 *versus* LIG3α at the downstream steps of the BER pathway. We showed the mutagenic ligation of resulting nick repair product by both BER ligases after polβ dATP:8-oxoG insertion, while there was a drastic decrease in the nick sealing efficiency by LIG3α after polβ inserts dCTP:8-oxoG, demonstrating that ambiguous coding potential and the identity of BER ligase modulate the fidelity of final steps. Furthermore, we observed the same ligation efficiency after insertion of both mismatches opposite 8-oxoG by polβ cancer-associated variant E288K. However, upon mutation of polβ active site residues K280 and R283 that play critical roles for discrimination between -*anti* and -*syn* conformations of 8-oxoG, our results revealed that the active site contacts govern nick sealing of mutagenic or error-free bypass of the lesion. Notably, the repair pathway coordination between polβ and BER ligases is dramatically diminished in the presence of ribonucleotide mismatches. Furthermore, in the ligation assays including DNA ligase alone, our results showed mutagenic end joining of 3’- dA:8-oxoG by LIG1 and LIG3α, while there was a completely diminished ligation of nick substrate with 3’-dC:8-oxoG by LIG3α. However, in the presence of a ribonucleotide at 3’-end, both BER ligases were able to seal RNA/DNA nicks, 3’-rA:8-oxoG and 3’-rC:8-oxoG, at similar efficiency. Finally, our results showed that APE1 removes 3’-ribonucleotides (rA or rC) more efficiently than 3’-mismatches (dA or dC) when paired with template 8-oxoG.

Overall, our findings demonstrate the deviations from canonical repair pathway coordination after polβ dNTP or rNTP mismatch incorporations during 8-oxoG bypass and how the dual coding potential of the oxidative lesion, in -*anti* versus -*syn* conformation, could govern mutagenic *versus* error-free outcomes during polβ gap filling and subsequent nick sealing by BER ligases. Our results also highlight the importance of functional interplay between APE1, polβ, LIG1, and LIG3α at the downstream steps in ensuring effective repair.

## Methods

### Protein purifications

DNA polymerase (pol) β (pET-28a) protein (1-335 amino acids) was purified as described (41–46). Briefly, the protein was overexpressed in BL21(DE3) *E. coli* cells in Lysogeny Broth (LB) media at 37 °C for 8 h and induced with 0.5 mM isopropyl β-D-thiogalactoside (IPTG). The cells were then grown overnight at 16 °C. After cell lysis at 4 °C by sonication in the lysis buffer containing 1X PBS (pH 7.3), 200 mM NaCl, 1 mM Dithiothreitol (DTT), and complete protease inhibitor cocktail, the lysate was pelleted at 16,000 x rpm for 1 h and then clarified by centrifugation and filtration. The supernatant was loaded on HisTrap HP column and purified with an increasing imidazole gradient (0-300 mM) elution at 4 °C. The collected fractions including his-tag polβ protein were then subsequently loaded on HiTrap Heparin column and eluted with a linear gradient of NaCl up to 1 M. Polβ active site mutants R283K and K280A as well as polβ cancer-associated variant E288K were purified as described for wild-type enzyme above. DNA ligase (LIG) 3α (pET-24b) full-length (1-922 amino acids) protein was purified as described (41–46). Briefly, the protein was overexpressed in BL21(DE3) *E. coli* cells in LB media at 37 °C for 8 h and induced with 0.5 mM IPTG. The cells were harvested, lysed at 4 °C, and then clarified as described above. The supernatant was loaded onto HisTrap HP column and purified with an increasing imidazole gradient (0-300 mM) elution at 4 °C. The collected fractions including his-tag LIG3α protein were then further purified by Heparin with a linear gradient of NaCl up to 1 M and then finally by Superdex 200 Increase 10/300 column in the buffer containing 50 mM Tris-HCl (pH 7.0), 500 mM NaCl, 5% glycerol, and 1 mM DTT. DNA ligase (LIG) 1 (pET-24b) full-length (1-919 amino acids) protein was purified as described (41–46). Briefly, the protein was overexpressed in BL21(DE3) *E. coli* cells and the cells were harvested, lysed at 4 °C, and then clarified as described above. The supernatant was loaded on HisTrap HP column and purified with an increasing imidazole gradient (0-300 mM) elution at 4 °C. The collected fractions were then subsequently loaded on HiTrap Heparin column with a linear gradient of NaCl up to 1 M. His-tag LIG1 protein was then further purified by Superdex 200 Increase 10/300 column in the buffer containing 50 mM Tris-HCl (pH 7.0), 500 mM NaCl, 5% glycerol, and 1 mM DTT. LIG1 deficiency disease-associated mutants P529L, R641L, and R771W were purified as described for wild-type enzyme above. AP-Endonuclease 1 (APE1) full-length (1-318 amino acids) protein (pET-24b) was purified as described (41–46). Briefly, the protein was overexpressed in BL21(DE3) *E. coli* cells and the cells were harvested, lysed at 4 °C, and the supernatant was loaded onto HisTrap HP column as described above. His-tag APE1 protein was purified with an increasing imidazole gradient (0-300 mM) elution and then loaded onto HiTrap Heparin column to further purify as a linear gradient of NaCl up to 1 M, and then finally loaded on Superdex 200 Increase 10/300 column in the buffer containing 50 mM Tris-HCl (pH 7.0), 500 mM NaCl, 5% glycerol, and 1 mM DTT. All proteins used in this study were dialyzed against storage buffer containing 25 mM TrisHCl (pH 7.4), 100 mM KCl, 1 mM TCEP, and 10% glycerol, concentrated, frozen in liquid nitrogen, and stored at -80 °C in aliquots. The final purity of all proteins is presented in Supplementary Figures S1-2.

### Polβ nucleotide insertion assays

Polβ nucleotide insertion assays were performed as described (41–46). One nucleotide gap DNA substrates with template 8-oxoG or C were used (Supplementary Table S1) to test polβ dNTP (dATP or dCTP) or rNTP (rATP or rCTP) insertion opposite 8-oxoG and correct dGTP insertion opposite C. The reaction mixture contains 50 mM Tris-HCl (pH 7.5), 100 mM KCl, 10 mM MgCl2, 1 mM ATP, 1 mM DTT, 100 µg ml^-1^ BSA, 1% glycerol, dNTP (100 µM), and DNA substrate (500 nM) in the final volume of 10 µl. The reaction was initiated by the addition of polβ (10 nM) and incubated at 37 °C for the time points as indicated in the figure legends. The reaction products were then mixed with an equal amount of gel loading buffer containing 95% formamide, 20 mM EDTA, 0.02% bromophenol blue, and 0.02% xylene cyanol and separated by electrophoresis on 18% Urea-PAGE gel. The gels were finally scanned with a Typhoon PhosphorImager (Amersham Typhoon RGB), and the data were analyzed using ImageQuant software. The nucleotide insertion assays were performed similarly for polβ wild-type and mutants (K280A, R283K, and E288K).

### Polβ nucleotide insertion coupled to DNA ligation assays

Coupled assays were performed as described (41–46). One nucleotide gap DNA substrates with template 8-oxoG or C (Supplementary Table S2) were used to test the ligation efficiency after polβ dNTP (dATP or dCTP) or rNTP (rATP or rCTP) insertion opposite 8-oxoG and polβ correct dGTP insertion opposite C by LIG1 and LIG3α. The reaction mixture contains 50 mM Tris-HCl (pH 7.5), 100 mM KCl, 10 mM MgCl2, 1 mM ATP, 1 mM DTT, 100 µg ml^-1^ BSA, 1% glycerol, dNTP (100 µM), and DNA substrate (500 nM) in the final volume of 10 µl. The reaction was initiated by the addition of polβ and DNA ligase protein complex (100 nM) and incubated at 37 °C for the time points as indicated in the figure legends. The reaction products were then mixed with an equal amount of gel loading buffer, separated by electrophoresis on 18% Urea-PAGE gel, and the data were analyzed using ImageQuant software as described above. The coupled assays were performed similarly for polβ wild-type and mutants (K280A, R283K, and E288K).

### DNA ligation assays

DNA ligation assays were performed as described (41–46). Nick DNA substrates with preinserted mismatches 3’-dA:8-oxoG or 3’-dC:8-oxoG and 3’-rA:8-oxoG or 3’-rC:8-oxoG as well as canonical 3’-dG:C (Supplementary Table S3) were used to test the nick sealing efficiency of LIG1 and LIG3α. The reaction mixture contains 50 mM Tris-HCl (pH 7.5), 100 mM KCl, 10 mM MgCl2, 1 mM ATP, 1 mM DTT, 100 µg ml^-1^ BSA, 1% glycerol, and DNA substrate (500 nM) in the final volume of 10 µl. The reaction was initiated by the addition of LIG1 or LIG3α (100 nM) and incubated at 37 °C for the time points as indicated in the figure legends. The reaction products were then mixed with an equal amount of gel loading buffer, separated by electrophoresis on 18% Urea-PAGE gel, and the data were analyzed using ImageQuant software as described above. The ligation assays were performed similarly for LIG1 deficiency disease-associated mutants (P529L, R641L, and R771W).

### APE1 exonuclease assays

APE1 exonuclease assays were performed as described (41–46). Nick DNA substrates with preinserted 3’-dA:8-oxoG or 3’-dC:8-oxoG and 3’-rA:8-oxoG or 3’-rC:8-oxoG were used (Supplementary Table S4) to test APE1 exonuclease activity for the removal of 3’-mismatched base (dA/rA or dC/rC) from template 8-oxoG. The reaction mixture contains 50 mM Tris-HCl (pH 7.5), 100 mM KCl, 10 mM MgCl2, 1 mM ATP, 1 mM DTT, 100 µg ml^-1^ BSA, 1% glycerol, and DNA substrate (500 nM) in the final volume of 10 µl. The reaction was initiated by the addition of APE1 alone (1 µM) and incubated at 37 °C for the time points as indicated in the figure legends. The reaction products were then mixed with an equal amount of gel loading buffer, separated by electrophoresis on 18% Urea-PAGE gel, and the data were analyzed using ImageQuant software as described above.

### APE1 exonuclease coupled to DNA ligation assays

Coupled assays were performed as described (41–46). Nick DNA substrates with preinserted 3’-dA:8-oxoG or 3’-dC:8-oxoG and 3’-rA:8-oxoG or 3’-rC:8-oxoG were used (Supplementary Table S5) to test ligation of nick by LIG1 or LIG3α coupled to the removal of 3’-mismatched base (dA/rA or dC/rC) from nick containing template 8-oxoG by APE1. The reaction mixture contains 50 mM Tris-HCl (pH 7.5), 100 mM KCl, 10 mM MgCl2, 1 mM ATP, 1 mM DTT, 100 µg ml^-1^ BSA, 1% glycerol, and DNA substrate (500 nM) in the final volume of 10 µl. The reaction was initiated by the addition of APE1 and DNA ligase protein complex (100 nM) and incubated at 37 °C for the time points as indicated in the figure legends. The reaction products were then mixed with an equal amount of gel loading buffer, separated by electrophoresis on 18% Urea-PAGE gel, and the data were analyzed using ImageQuant software as described above.

## Results

### Ligation of polβ mismatch insertion products during 8-oxoG bypass by LIG1 and LIG3α

We first investigated dATP and dCTP mismatch insertion by polβ during 8-oxoG bypass and subsequent nick sealing by LIG1 or LIG3α (Figure 1A). Polβ can insert dATP:8-oxoG and dCTP:8-oxoG at similar efficiency (Figure 1B and Supplementary Figure S3A). We then compared the ligation efficiency of these polβ mismatch insertion products by the BER ligases. In the presence of LIG1, the ligation products after both insertions were obtained (Figure 1C).

**Figure 1.**
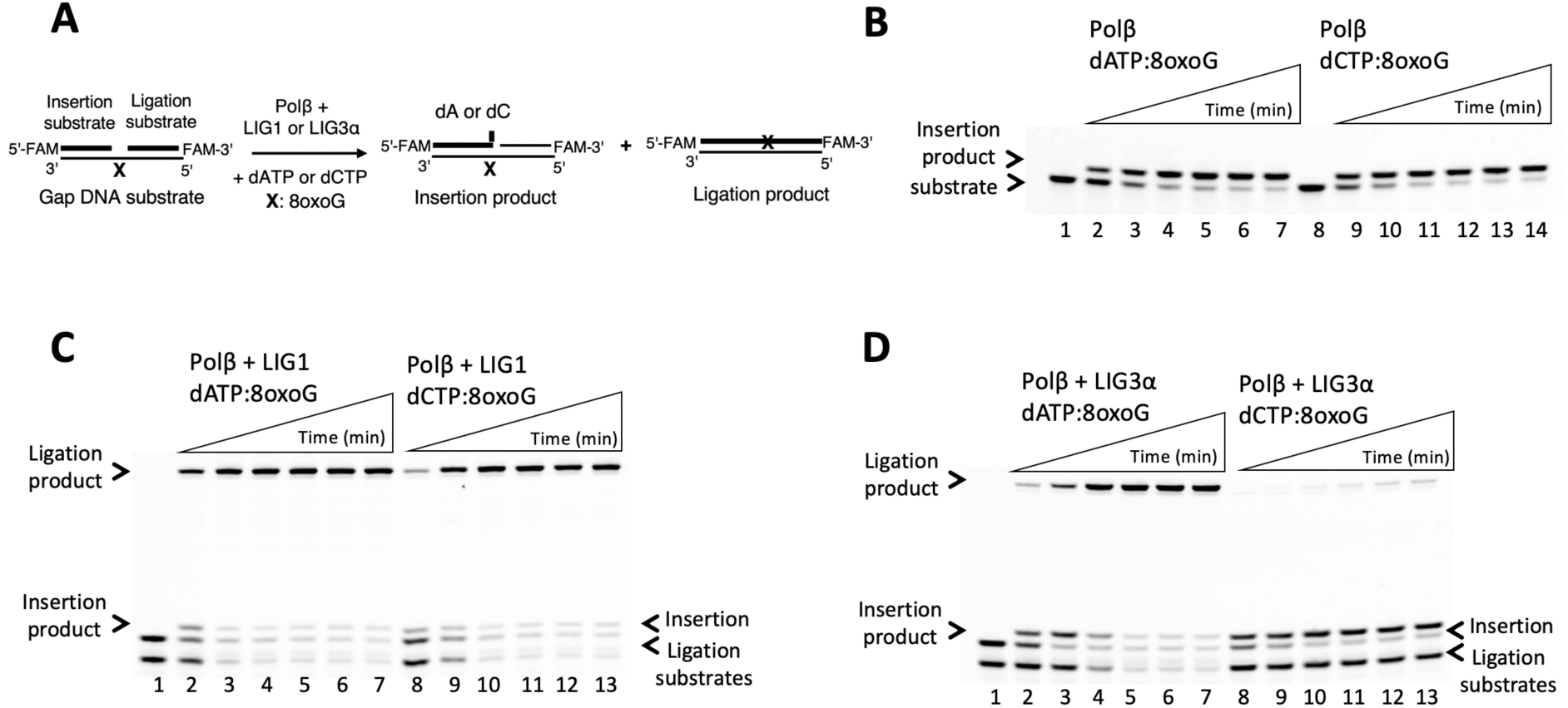
Ligation of polβ 8-oxoG bypass products by BER ligases. **(A)** Scheme shows gap substrate and reaction products in the coupled assay including polβ, dATP or dCTP, and BER ligases. **(B)** Lanes 1 and 8 are the negative enzyme controls of the one nucleotide gap DNA substrate with template 8-oxoG. Lanes 2-7 and 9-14 are polβ insertion products of dATP and dCTP opposite template 8-oxoG, respectively, and correspond to time points of 0.5, 1, 2, 3, 4, and 5 min. **(C-D)** Line 1 is the negative enzyme control of the one nucleotide gap DNA substrate with template 8-oxoG. Lanes 2-7 and 8-13 are the ligation products of polβ dATP and dCTP insertions, respectively, by LIG1 (C) and LIG3α (D), and correspond to time points of 0.5, 1, 2, 3, 4, and 5 min.

However, LIG3α was only able to seal resulting nick product of polβ dATP:8-oxoG insertion (Figure 1D, lanes 2-7), which was relatively less efficient when compared to that of LIG1, particularly at earlier time points of reaction (Figure 1C *versus* 1D, lanes 2-4). Interestingly, no ligation product was observed after polβ dCTP:8-oxoG insertion by LIG3α (Figure 1D, lanes 8-13), when compared to an efficient ligation by LIG1 (Figure 1C *versus* 1D, lanes 9- 13).

When comparing the ligation efficiency after polβ dATP:8-oxoG *versus* dCTP:8-oxoG insertions by the BER ligases, our results demonstrated that the nick sealing by LIG1 after both insertions showed a similar amount of ligation products (Figure 2A). However, this ligation efficiency is significantly reduced by LIG3α in case of polβ dCTP:8-oxoG insertion (Figure 2B). Similarly, the comparison of ligation efficiencies by LIG1 *versus* LIG3α depending on the type of mismatch inserted by polβ during 8-oxoG bypass showed similar nick sealing profile for dATP:8-oxoG (Figure 2C), while there was ∼80-fold more efficient nick sealing by LIG1 over LIG3α after polβ dCTP:8-oxoG insertion (Figure 2D). In the control experiments, we confirmed the ligation of polβ correct dGTP:C insertion products (Supplementary Figure S4A).

**Figure 2.**
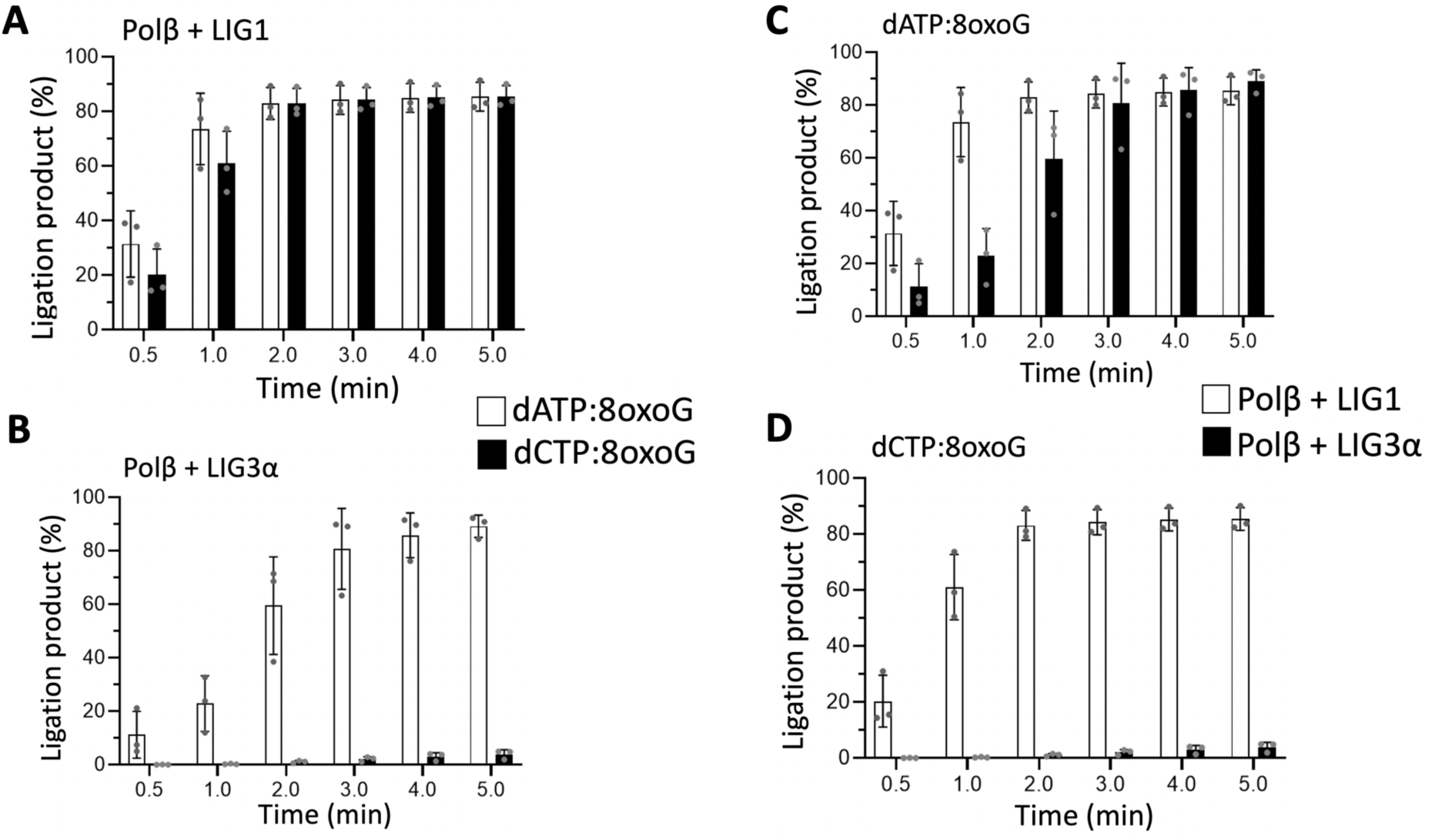
Comparison of ligation efficiency after polβ mismatch insertions during 8-oxoG bypass by LIG1 and LIG3α. (A-D) Graphs show time-dependent changes in the amount of ligation products by LIG1 and LIG3α after polβ dATP and dCTP insertions opposite template 8-oxoG. The data represent the average of three independent experiments ± SD.

### Impact of polβ active site mutations on the ligation efficiency of 8-oxoG bypass products

The X-ray structures of polβ binary complexes with an incoming dATP and dCTP opposite template 8-oxoG of a gap DNA reported the importance of polβ active site residues Lys(K)280 and Arg(R)283 for stabilization of the mutagenic -*syn* and non-mutagenic -*anti* conformations of 8-oxoG base pairing at the template-primer terminus (34–39). To further characterize the repair pathway coordination during polβ 8-oxoG bypass in both conformations, and to investigate the impact of mutations at these critical active sites, we generated polβ K280A and R383K mutants and compared the ligation efficiency after mismatch insertions as described above for wild-type enzyme (Figures 3A and 4A).

**Figure 3.**
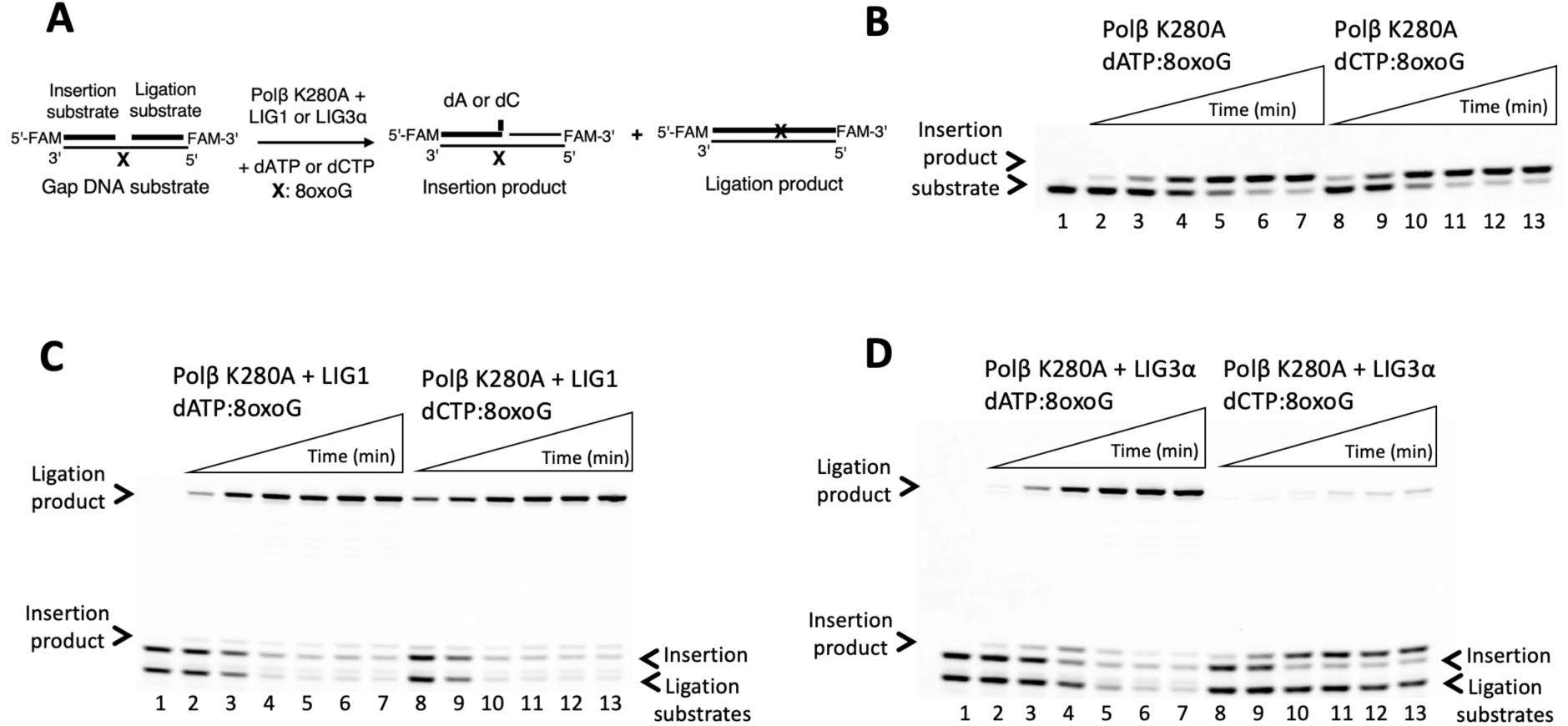
Ligation of polβ K280A 8-oxoG bypass products by BER ligases. **(A)** Scheme shows gap substrate and reaction products in the coupled assay including polβ K280A, dATP or dCTP, and BER ligases. **(B)** Line 1 is the negative enzyme control of the one nucleotide gap DNA substrate with template 8-oxoG. Lanes 2-7 and 8-13 are the insertion products of dATP and dCTP opposite template 8-oxoG, respectively, by polβ K280A mutant, and correspond to time points of 0.5, 1, 2, 3, 4, and 5 min. **(C-D)** Line 1 is the negative enzyme control of the one nucleotide gap DNA substrate with template 8-oxoG. Lanes 2-7 and 8-13 are the ligation products of polβ K280A dATP and dCTP insertions, respectively, by LIG1 (C) and LIG3α (D), and correspond to time points of 0.5, 1, 2, 3, 4, and 5 min.

**Figure 4.**
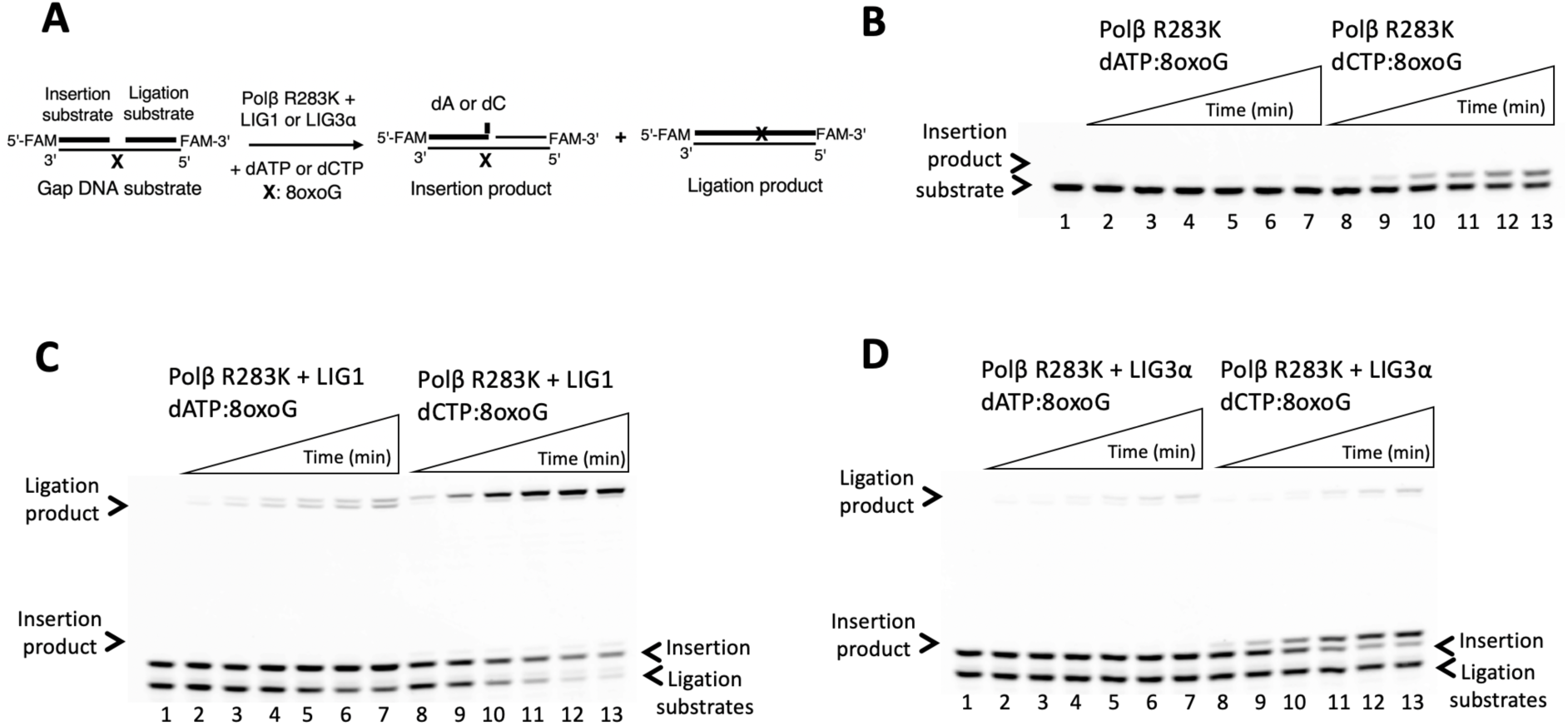
Ligation of polβ R283K 8-oxoG bypass products by BER ligases. **(A)** Scheme shows gap substrate and reaction products in the coupled assay including polβ R283K, dATP or dCTP, and BER ligases. **(B)** Line 1 is the negative enzyme control of the one nucleotide gap DNA substrate with template 8-oxoG. Lanes 2-7 and 8-13 are the insertion products of dATP and dCTP opposite template 8-oxoG, respectively, by polβ R283K mutant, and correspond to time points of 0.5, 1, 2, 3, 4, and 5 min. **(C-D)** Line 1 is the negative enzyme control of the one nucleotide gap DNA substrate with template 8-oxoG. Lanes 2-7 and 8-13 are the ligation products of polβ R283K dATP and dCTP insertions, respectively, by LIG1 (C) and LIG3α (D), and correspond to time points of 0.5, 1, 2, 3, 4, and 5 min.

While we observed dATP:8-oxoG and dCTP:8-oxoG insertions by polβ K280A (Figure 3B and Supplementary Figure S3B), the efficiency of dCTP:8-oxoG insertion was significantly reduced by the effect of R283K mutation that was not able to insert dATP:8-oxoG (Figure 4B and Supplementary Figure S3C). In the presence of polβ K280A and LIG1, we observed the ligation of dATP:8-oxoG and dCTP:8-oxoG insertion products at similar efficiency (Figure 3C). In the presence of polβ K280A and LIG3α, this efficiency for converting dATP:8-oxoG insertion to ligation products was slightly lower than that of LIG1 (Figure 3D, lanes 2-7) and no ligation product was observed following dCTP:8-oxoG insertion by polβ K280A (Figure 3D, lanes 8-13). However, R283K mutation adversely impacts the ligation after mismatch insertions (Figure 4C-D). We only observed nick sealing by LIG1 after dCTP:8-oxoG insertion (Figure 4C, lanes 8-13).

Our results demonstrated wild-type level of ligation efficiency by both BER ligases after mismatch insertions by polβ K280A mutant (Figure 5A-B). For R283K, there was a time-dependent increase in ligation product after dCTP:8-oxoG insertion by LIG1 (Figure 5C). Conversely, the ligation efficiency of LIG3α was significantly reduced when compared to LIG1 for both insertions by polβ R283K, showing a ∼60-fold difference (Figure 5D). Overall comparisons showed that the mutations at the polβ active sites affects gap filling, and therefore, subsequent nick sealing efficiency depending on a mismatch being inserted in *anti-* vs *syn-* conformation during 8-oxoG bypass (Supplementary Figure S5). In the control experiments, we observed differences in the ligation efficiency after a correct dGTP:C insertion by polβ active site mutants K280A and R283K (Supplementary Figure S4B-C).

**Figure 5.**
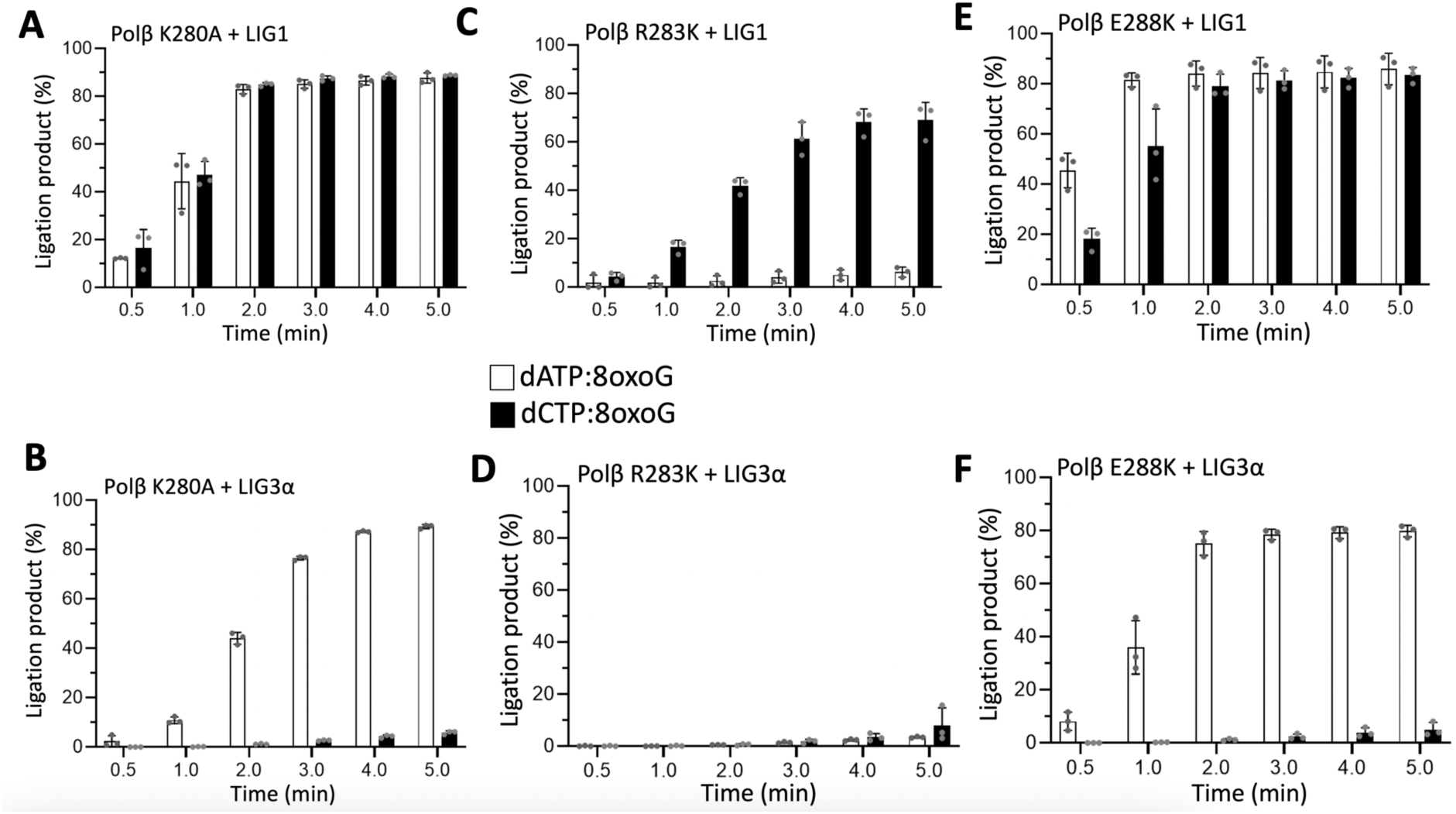
Comparison of ligation efficiency by LIG1 and LIG3α during 8-oxoG bypass by polβ active site mutants and cancer-associated variant. (A-F) Graphs show time- dependent changes in the amount of ligation products by LIG1 and LIG3α after dATP and dCTP insertions opposite template 8-oxoG by polβ active site mutants K280A, R283K and cancer-associated variant E288K. The data represents the average of three independent experiments ± SD.

### Ligation of mismatch insertion products by polβ cancer-associated variant during 8-oxoG bypass

In addition to the polβ active site mutants, we also questioned how cancer-associated mutations that have been identified (30% of human tumors) in *POLB* gene could affect the efficiency of downstream BER steps during 8-oxoG bypass. Polβ E288K, a colon tumor variant, exhibits an enhanced mutagenesis due to loss of fidelity although it shows similar gap DNA binding affinity with wild-type enzyme (47,48). We investigated the ligation efficiency of mismatch insertions opposite template 8-oxoG by polβ E288K variant (Figure 6A).

**Figure 6.**
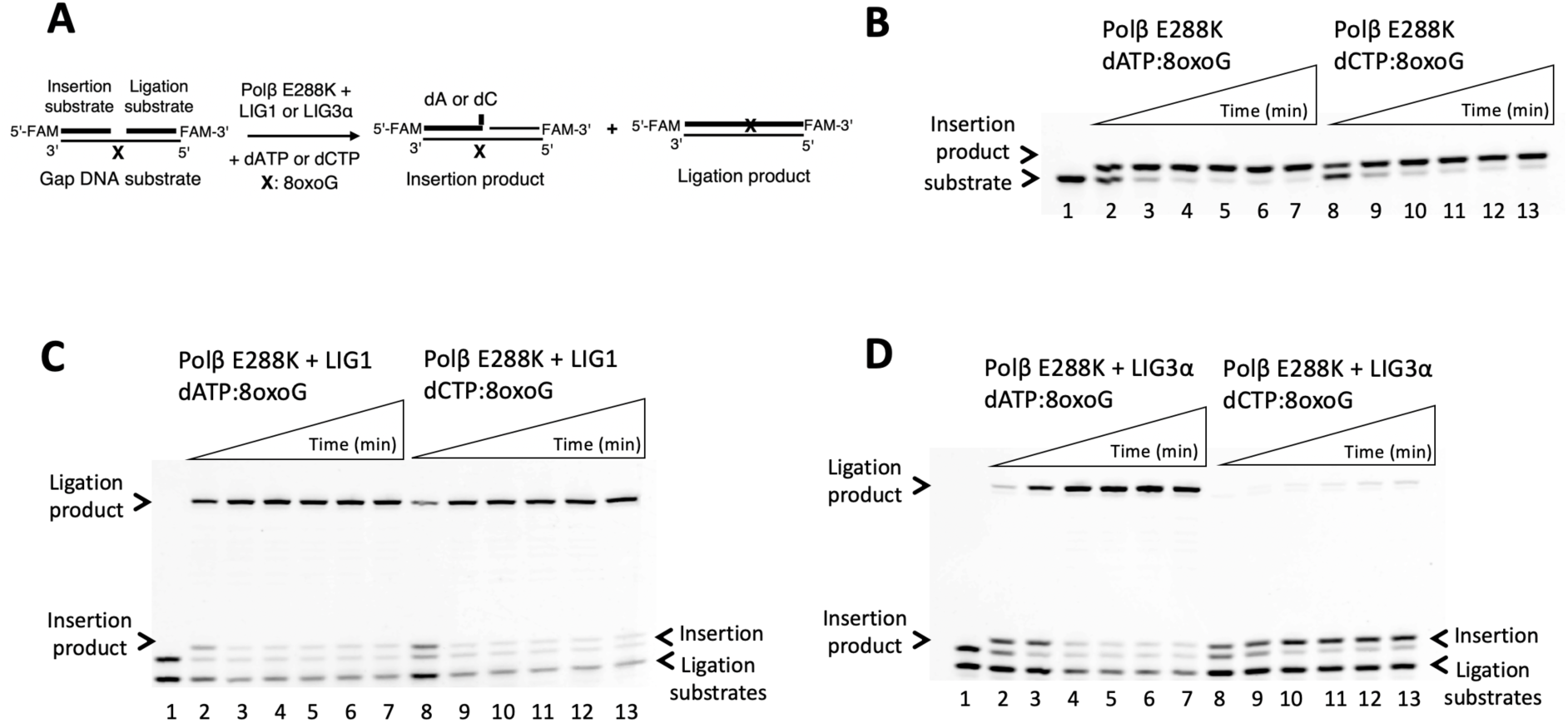
Ligation of polβ E288K 8-oxoG bypass products by BER ligases. **(A)** Scheme shows gap substrate and reaction products in the coupled assay including polβ E288K, dATP or dCTP, BER ligases. **(B)** Line 1 is the negative enzyme control of the one nucleotide gap DNA substrate with template 8-oxoG. Lanes 2-7 and 8-13 are the insertion products of dATP and dCTP opposite template 8-oxoG, respectively, by polβ E288K mutant, and correspond to time points of 0.5, 1, 2, 3, 4, and 5 min. **(C-D)** Line 1 is the negative enzyme control of the one nucleotide gap DNA substrate with template 8-oxoG. Lanes 2-7 and 8-13 are the ligation products of polβ E288K dATP and dCTP insertions, respectively, by LIG1 (C) and LIG3α (D), and correspond to time points of 0.5, 1, 2, 3, 4, and 5 min.

Polβ E288K can insert dATP:8-oxoG and dCTP:8-oxoG at similar efficiency (Figure 6B and Supplementary Figure 3D). In the presence of LIG1, we observed an efficient ligation after both insertions (Figure 6C). This ligation efficiency was relatively less in the presence of LIG3α after dATP:8-oxoG insertion (Figure 6D, lanes 2-7) and no ligation product was observed after dCTP:8-oxoG insertion (Figure 6D, lanes 8-13). Overall, we found that LIG1 shows similar efficiency in sealing nick products after both insertions by polβ E288K variant while the ligation of dCTP:8-oxoG insertion products by LIG3α was significantly reduced (Figure 5E-F). These results were found to be similar with polβ wild-type (Supplementary Figures S7-8).

### Ligation efficiency of nick DNA containing 3**’**-mismatch and template 8-oxoG by LIG1 and LIG3α

In addition to polβ mismatch insertion coupled to DNA ligation assays, to further investigate the efficiency of final steps in the presence of template 8-oxoG, we investigated the end joining ability of the BER ligases in ligation assays including nick DNA substrates with preinserted 3’- dA and 3’-dC mismatches opposite 8-oxoG by LIG1 wild-type, LIG1 deficiency disease-associated variants, and LIG3α (Figures 7A and 9A).

**Figure 7.**
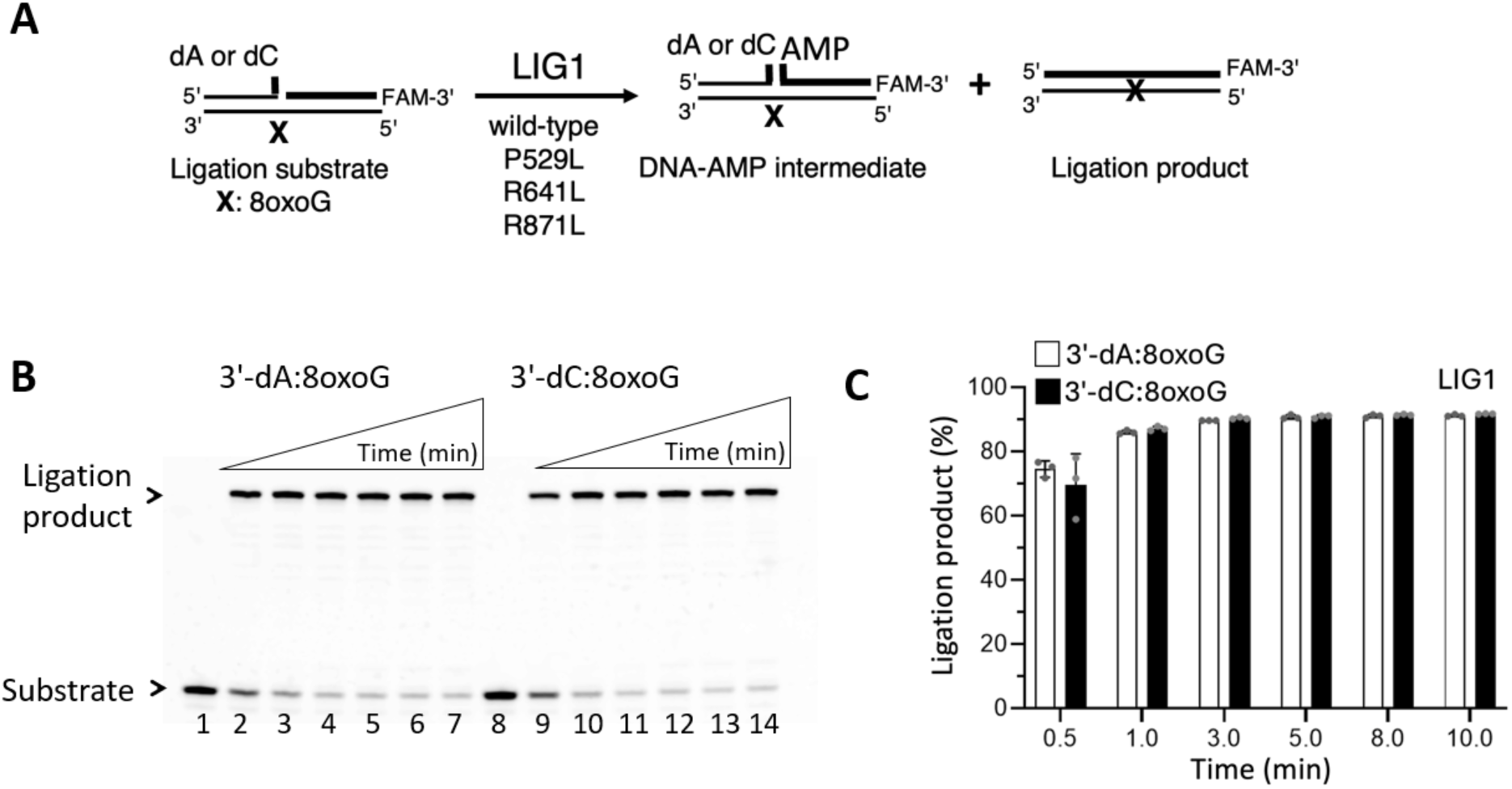
Ligation of nick DNA with template 8-oxoG by LIG1. **(A)** Scheme shows nick DNA substrate and reaction products in the ligation assay including LIG1 wild-type or variants. **(B)** Lanes 1 and 8 are the negative enzyme controls of the nick DNA substrates 3’-dA:8-oxoG and 3’-dC:8-oxoG, respectively. Lanes 2-7 and 9-14 are the ligation products by LIG1 in the presence of the nick DNA substrates 3’-dA:8-oxoG and 3’-dC:8-oxoG, respectively, and correspond to time points of 0.5, 1, 3, 5, 8, and 10 min. **(C)** Graph shows time-dependent changes in the amount of ligation products and the data represent the average of three independent experiments ± SD.

Our results showed that LIG1 can indiscriminately ligate nick DNA substrates containing 3’- dA:8-oxoG and 3’-dC:8-oxoG (Figure 7B). The amount of ligation products was similar despite base-pairing architecture (Figure 7C). We then questioned the impact of LIG1 deficiency disease-associated mutations, P529L, R641L, R771W on the ligation of nicks containing 8- oxoG on a template position (Figure 8). In the presence of LIG1 P529L variant, we observed similar ligation efficiency (Figure 8A), while LIG1 R641L showed significantly less ligation (Figure 8B) for both 3’-dA:8-oxoG and 3’-dC:8-oxoG substrates. There was a relatively less nick sealing of 3’-dC:8-oxoG by LIG1 R771W (Figure 8C). The amount of ligation products showed a time-dependent increase for all LIG1 variants, and the most dramatic decrease was observed with LIG1 R641L and R771W variants especially at earlier time points (Figures 8D- F). The comparison of ligation products between LIG1 wild-type *versus* variants also demonstrated the impact of R641L and R771W mutations on the ligation efficiency of 3’-dA:8- oxoG and 3’-dC:8-oxoG substrates (Supplementary Figure S9). In the control assays, we observed ligation of canonical nick by all LIG1 mutants (Supplementary Figure S10).

**Figure 8.**
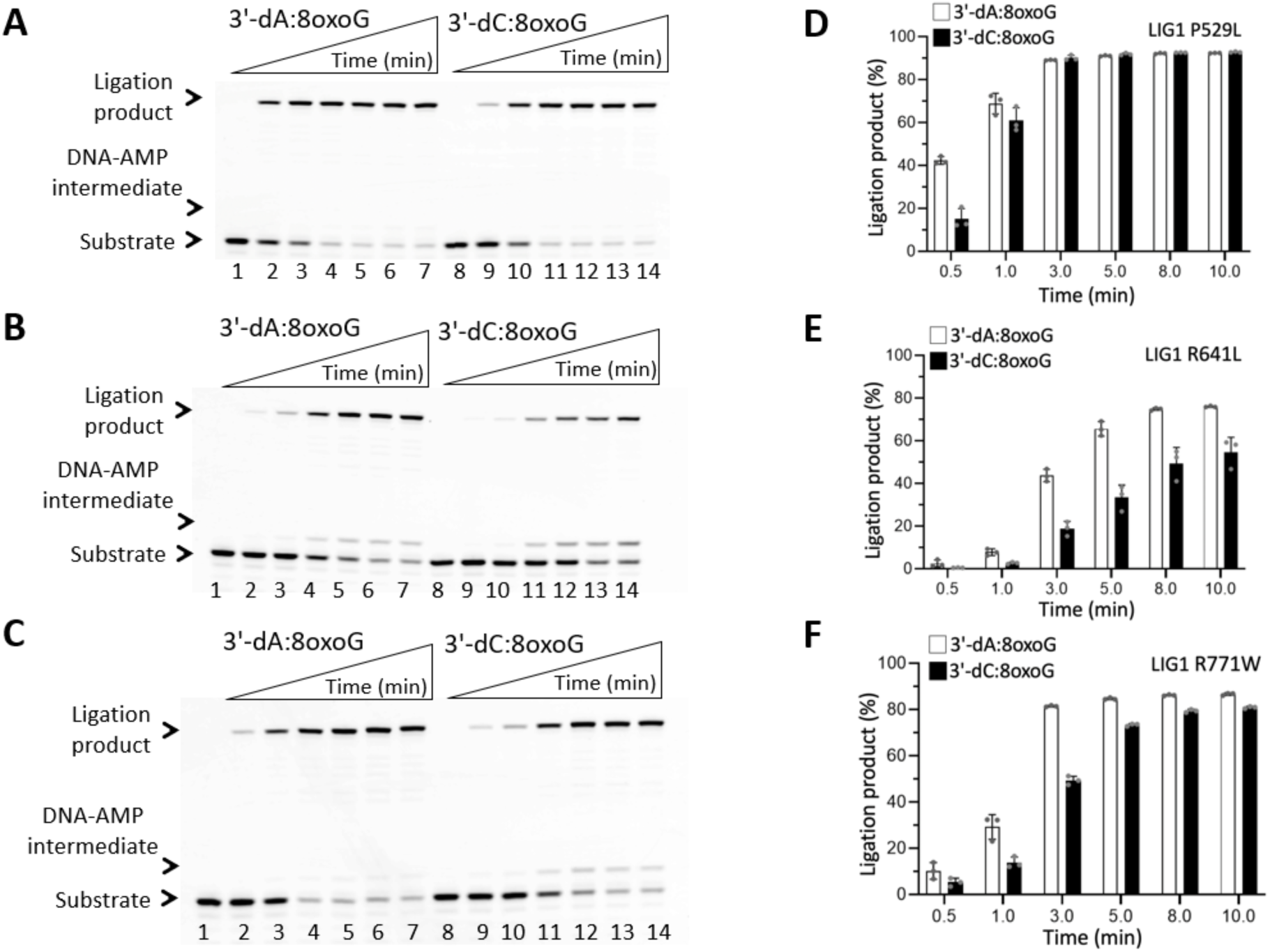
Ligation of nick DNA with template 8-oxoG by LIG1 variants. (A-C) Lanes 1 and 8 are the negative enzyme controls of the nick DNA substrates 3’-dA:8-oxoG and 3’-dC:8- oxoG, respectively. Lanes 2-7 and 9-14 are the ligation products by LIG1 deficiency disease- associated variants P529L (A), R641L (B), and R771W (C) in the presence of the nick DNA substrates 3’-dA:8-oxoG and 3’-dC:8-oxoG, respectively, and correspond to time points of 0.5, 1, 3, 5, 8, and 10 min. **(D-F)** Graphs show time-dependent changes in the amount of ligation products and the data represent the average of three independent experiments ± SD.

Regarding LIG3α (Figure 9A), our results showed an efficient ligation only in the presence of 3’-dA:8-oxoG (Figure 9B-C). The comparison of ligation efficiency by LIG1 *versus* LIG3α showed difference depending on the nick substrate. LIG1 exhibits similar efficiency for both nick substrates, and we observed ∼90-fold difference in the amount of ligation product for 3’- dC:8-oxoG substrate by LIG3α (Figure 9D-E). In the control assays, we observed an efficient ligation of canonical nick by both BER ligases (Supplementary Figure S11).

**Figure 9.**
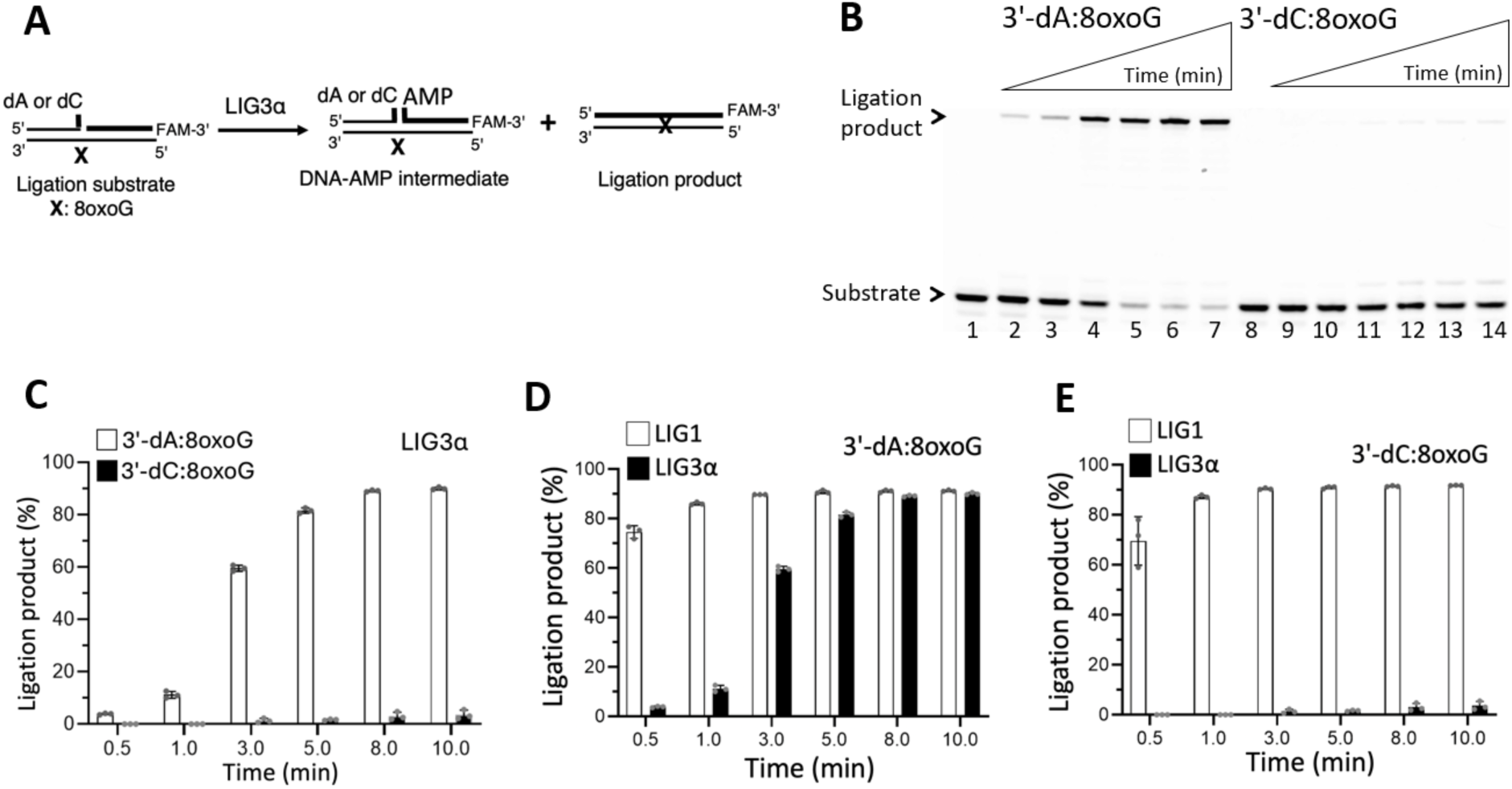
Ligation of nick DNA with template 8-oxoG by LIG3α. **(A)** Scheme shows nick DNA substrate and reaction products in the ligation assay including LIG3α. **(B)** Lanes 1 and 8 are the negative enzyme controls of the nick DNA substrates with 3’-dA:8-oxoG and 3’-dC:8- oxoG, respectively. Lanes 2-7 and 9-14 are the ligation products in the presence of the nick DNA substrates 3’-dA:8-oxoG and 3’-dC:8-oxoG, respectively, by LIG3α, and correspond to time points of 0.5, 1, 3, 5, 8, and 10 min. (**C-E**) Graphs show time-dependent changes in the amount of ligation products and the data represent the average of three independent experiments ± SD.

### Impact of polβ ribonucleotide incorporation during 8-oxoG bypass on the efficiency of nick sealing by BER ligases

To comprehensively elucidate the impact of mismatch insertions by polβ bypassing 8-oxoG on the repair pathway coordination with the BER ligases, we then questioned how rNTP incorporation opposite 8-oxoG could affect the next nick sealing step (Figure 10A). Polβ fails to insert rATP:8-oxoG and rCTP:8-oxoG (Figure 10B and Supplementary Figure S3E). In the presence of LIG1 or LIG3α, our results demonstrated no ligation at all (Figure 10C-D). We only obtained a gap ligation product by LIG1, which refers to the ligation of one nucleotide gap substrate itself as shown by the difference in size with a complete ligation product after polβ dATP:8-oxoG (Figure 10C, line 2 *versus* lanes 3-8) or dCTP:8-oxoG (Figure 10C, line 9 *versus* lanes 10-15) insertions. These results demonstrated that the nick sealing efficiency after polβ dNTP *versus* rNTP mismatch insertions is drastically different (Figure 11), suggesting that 8-oxoG bypass by polβ is dictated by the nature of an incoming nucleotide and identity of the sugar, and thereby, promoting insertion of dATP in *syn*-8-oxoG conformation by polβ primarily leads to mutagenic ligation of resulting nick product at the final steps of the BER pathway.

**Figure 10.**
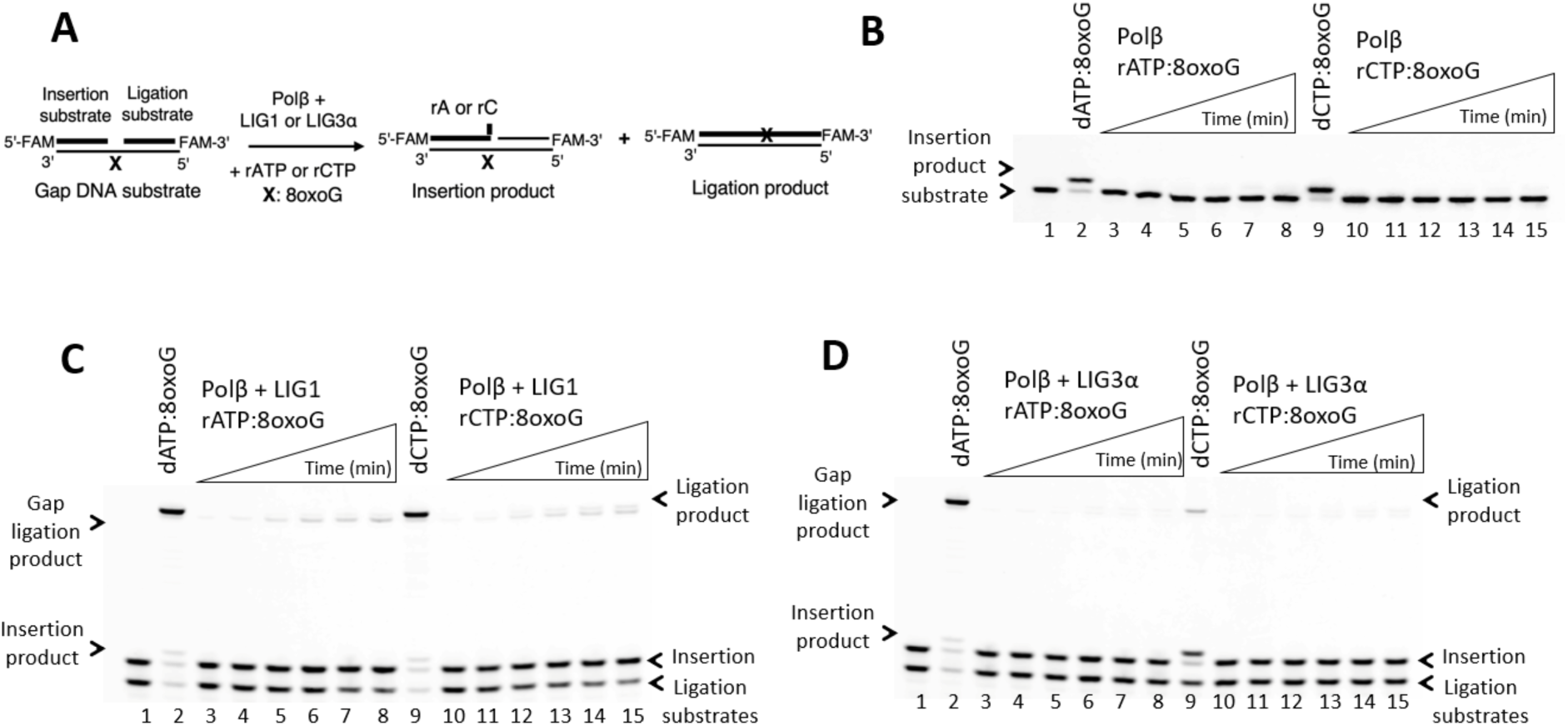
Ligation of polβ ribonucleotide insertion products during 8-oxoG bypass by LIG1 and LIG3α. **(A)** Scheme shows gap substrate and reaction products in the coupled assay including polβ, rATP or rCTP, and BER ligases. **(B)** Lanes 1 and 8 are the negative enzyme controls of the one nucleotide gap DNA substrate with template 8-oxoG. Lanes 2 and 9 are polβ insertion products of rATP and rCTP opposite template 8-oxoG, respectively, and correspond to time points of 0.5, 1, 2, 3, 4, and 5 min. Lanes 3-8 and 10-15 are polβ insertion products of rATP and rCTP opposite template 8-oxoG, respectively, and correspond to time points of 0.5, 1, 2, 3, 4, and 5 min. **(C-D)** Lane 1 is the negative enzyme control of the one nucleotide gap DNA substrate with template 8-oxoG. Lanes 2 and 9 are ligation products of polβ rATP and rCTP insertions by LIG1 (C) and LIG3α (D). Lanes 3-8 and 10-15 are the ligation products of polβ rATP and rCTP insertions, respectively, by LIG1 (C) and LIG3α (D), and correspond to time points of 0.5, 1, 2, 3, 4, and 5 min.

**Figure 11.**
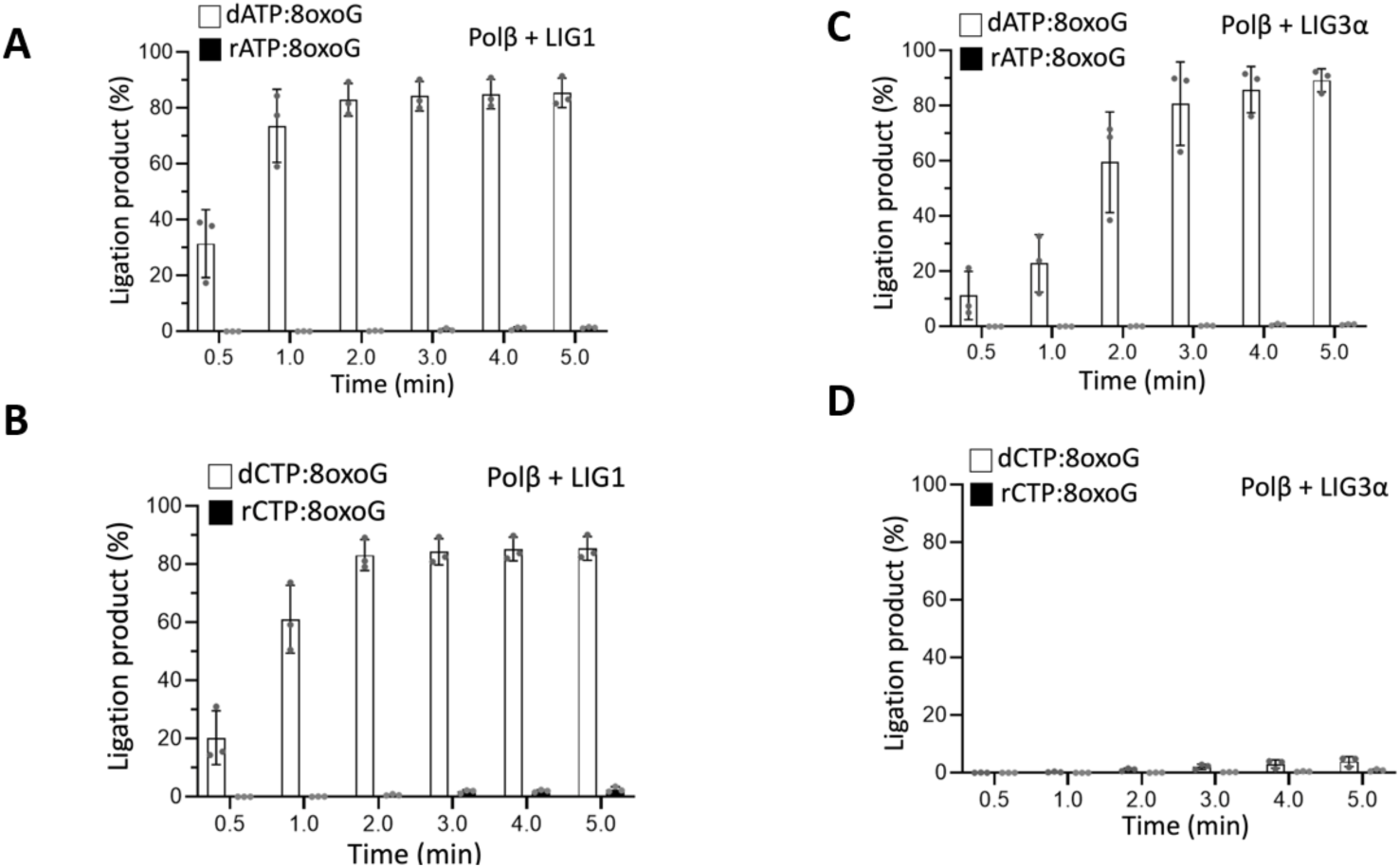
Comparison of ligation efficiency after polβ mismatch *versus* ribonucleotide insertion during 8-oxoG bypass by LIG1 and LIG3α. (A-D) Graphs show time-dependent changes in the amount of ligation products by LIG1 and LIG3α after polβ dATP vs rATP and dCTP vs rCTP insertions opposite template 8-oxoG. The data represent the average of three independent experiments ± SD.

### Ligation efficiency of nick DNA containing 3**’**-ribonucleotide and template 8-oxoG by LIG1 and LIG3α

We also tested how the presence of a single ribonucleotide at the 3’-end opposite template 8- oxoG on nick DNA affects the end joining abilities of LIG1 and LIG3α (Figures 12A and 14A). LIG1 can efficiently seal both nick substrates 3’-rA:8-oxoG and 3’-rC:8-oxoG (Figure 12B-C). Similarly, LIG1 deficiency disease-associated variants P529L, R641L, and R771W can join DNA ends with 3’-rA and 3’-rC opposite template 8-oxoG (Figure 13). When compared to wild-type enzyme, we observed a decrease in nick sealing efficiency of 3’-rA:8-oxoG and 3’- rC:8-oxoG substrates at earlier time points of ligation reaction by LIG1 variants (Supplementary Figure S12).

**Figure 12.**
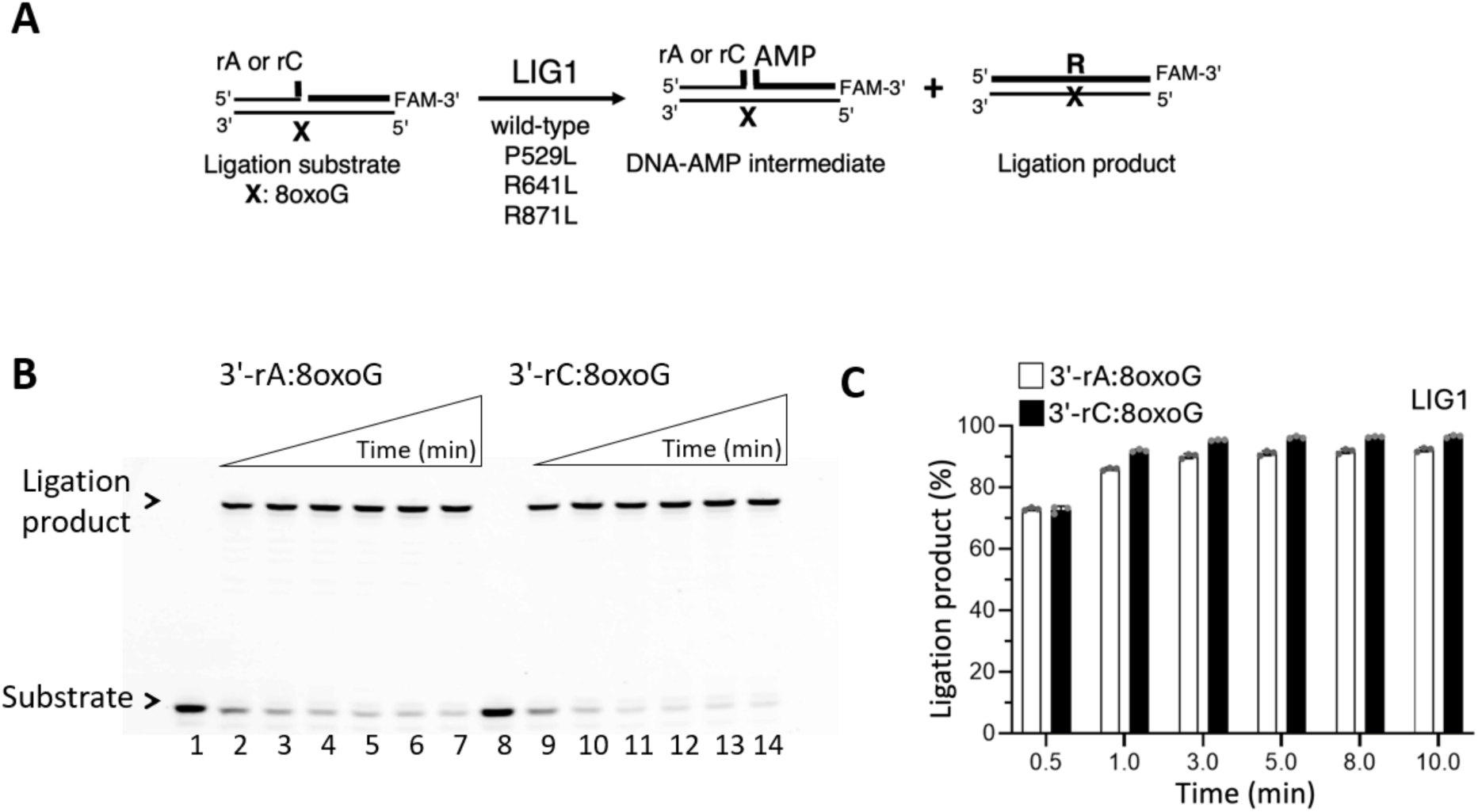
Ligation of nick DNA with 3’-ribonucleotide and template 8-oxoG by LIG1. **(A)** Scheme shows nick DNA substrates with 3’-ribonucleotide opposite 8-oxoG and reaction products in the ligation assay including LIG1. **(B)** Lanes 1 and 8 are the negative enzyme controls of the nick DNA substrates 3’-rA:8-oxoG and 3’-rC:8-oxoG, respectively. Lanes 2-7 and 9-14 are the ligation products by LIG1 in the presence of the nick DNA substrates 3’-rA:8- oxoG and 3’-rC:8-oxoG, respectively, and correspond to time points of 0.5, 1, 3, 5, 8, and 10 min. **(C)** Graph shows time-dependent changes in the amount of ligation products and the data represent the average of three independent experiments ± SD.

**Figure 13.**
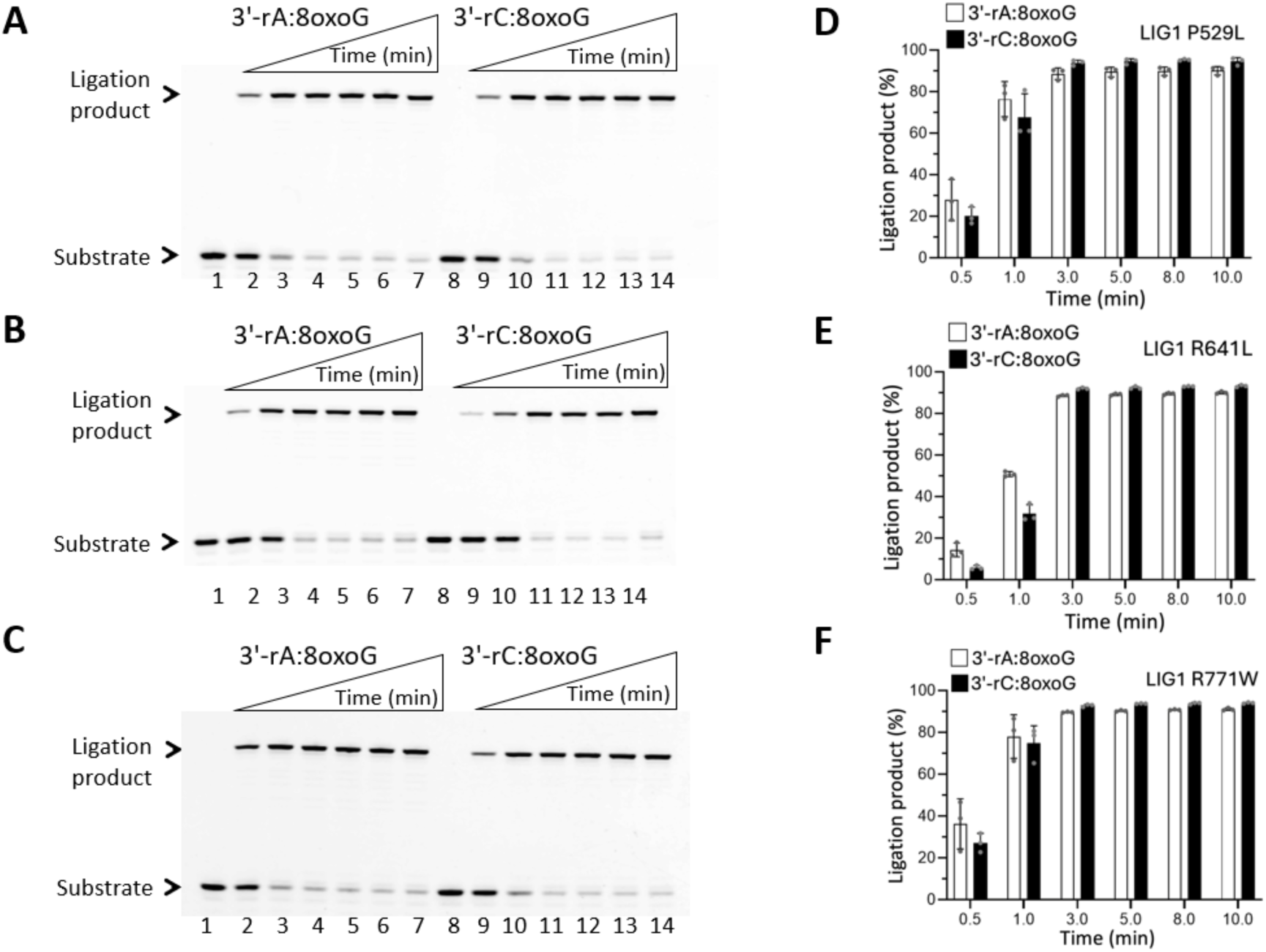
Ligation of nick DNA with 3’-ribonucleaotide and template 8-oxoG by LIG1 variants. (A-C) Lanes 1 and 8 are the negative enzyme controls of the nick DNA substrates 3’-rA:8-oxoG and 3’-rC:8-oxoG, respectively. Lanes 2-7 and 9-14 are the ligation products by LIG1 deficiency disease-associated variants P529L (A), R641L (B), and R771W (C) in the presence of the nick DNA substrates 3’-rA:8-oxoG and 3’-rC:8-oxoG, respectively, and correspond to time points of 0.5, 1, 3, 5, 8, and 10 min. **(D-F)** Graphs show time-dependent changes in the amount of ligation products and the data represent the average of three independent experiments ± SD.

**Figure 14.**
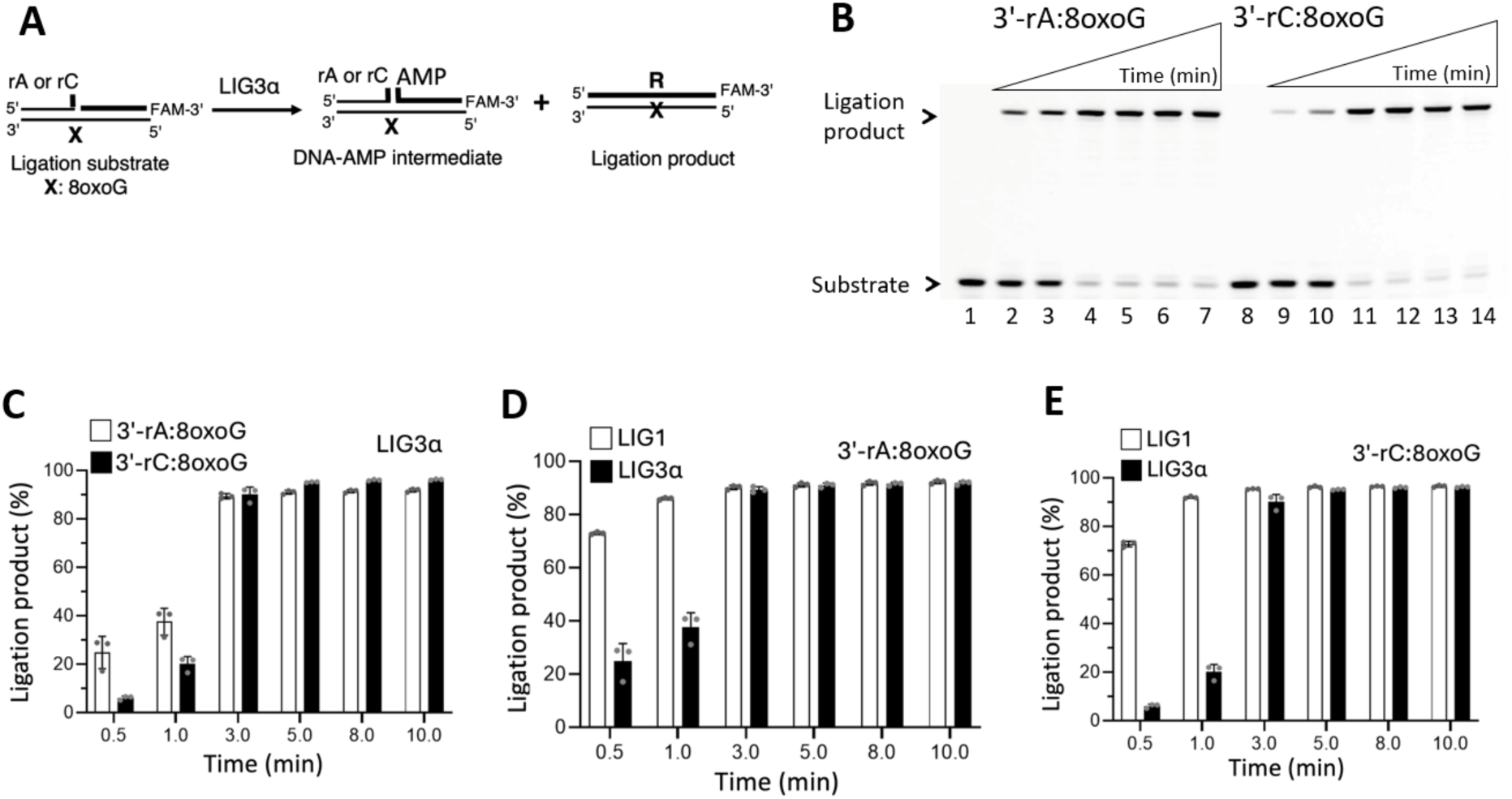
Ligation of nick DNA with 3’-ribonucleotide and template 8-oxoG by LIG3α. **(A)** Scheme shows nick DNA substrates with 3’-ribonucleotide opposite 8-oxoG and reaction products in the ligation assay including LIG3α. **(B)** Lanes 1 and 8 are the negative enzyme controls of the nick DNA substrates with 3’-rA:8-oxoG and 3’-rC:8-oxoG, respectively. Lanes 2-7 and 9-14 are the ligation products in the presence of the nick DNA substrates with 3’-rA:8- oxoG and 3’-rC:8-oxoG, respectively, by LIG3α, and correspond to time points of 0.5, 1, 3, 5, 8, and 10 min. (**C-E**) Graphs show time-dependent changes in the amount of ligation products and the data represent the average of three independent experiments ± SD.

Regarding LIG3α, our results showed a lack of sugar discrimination against 3’-ribo (rA or rC) when base paired with template 8-oxoG (Figure 14B) and the nick sealing of 3’-rC:8-oxoG was relatively less efficient than that of 3’-rA:8-oxoG (Figure 14C). The comparison of ligation products by the BER ligases demonstrated that LIG1 shows higher efficiency for both nick substrates, *i.e*, ∼6-fold difference in the amount of ligation product with 3’-rC:8-oxoG (Figure 14D-E and Supplementary Figure S13). Lastly, we compared how the identity of sugar (dA or rA *versus* dC or rC) opposite template 8-oxoG affects end joining ability of the BER ligases. LIG1 was able to ligate all nick substrates with similar efficiency. However, LIG3α can seal nicks with 3’-dA:8-oxoG, 3’-rA:8-oxoG, and 3’-rC:8-oxoG but is uanble to ligate nick with 3’- dC:8-oxoG (Supplementary Figure S14).

### Interplay between APE1 and BER ligases during the removal or ligation of 3’-mismatches from the nick repair intermediates containing template 8-oxoG

To comprehensively elucidate the molecular determinants of BER accuracy during polβ 8- oxoG bypass at the downstream steps, we finally investigated the proofreading activity of APE1 and its coordination with the BER ligases (Figures 15A and 16A).

**Figure 15.**
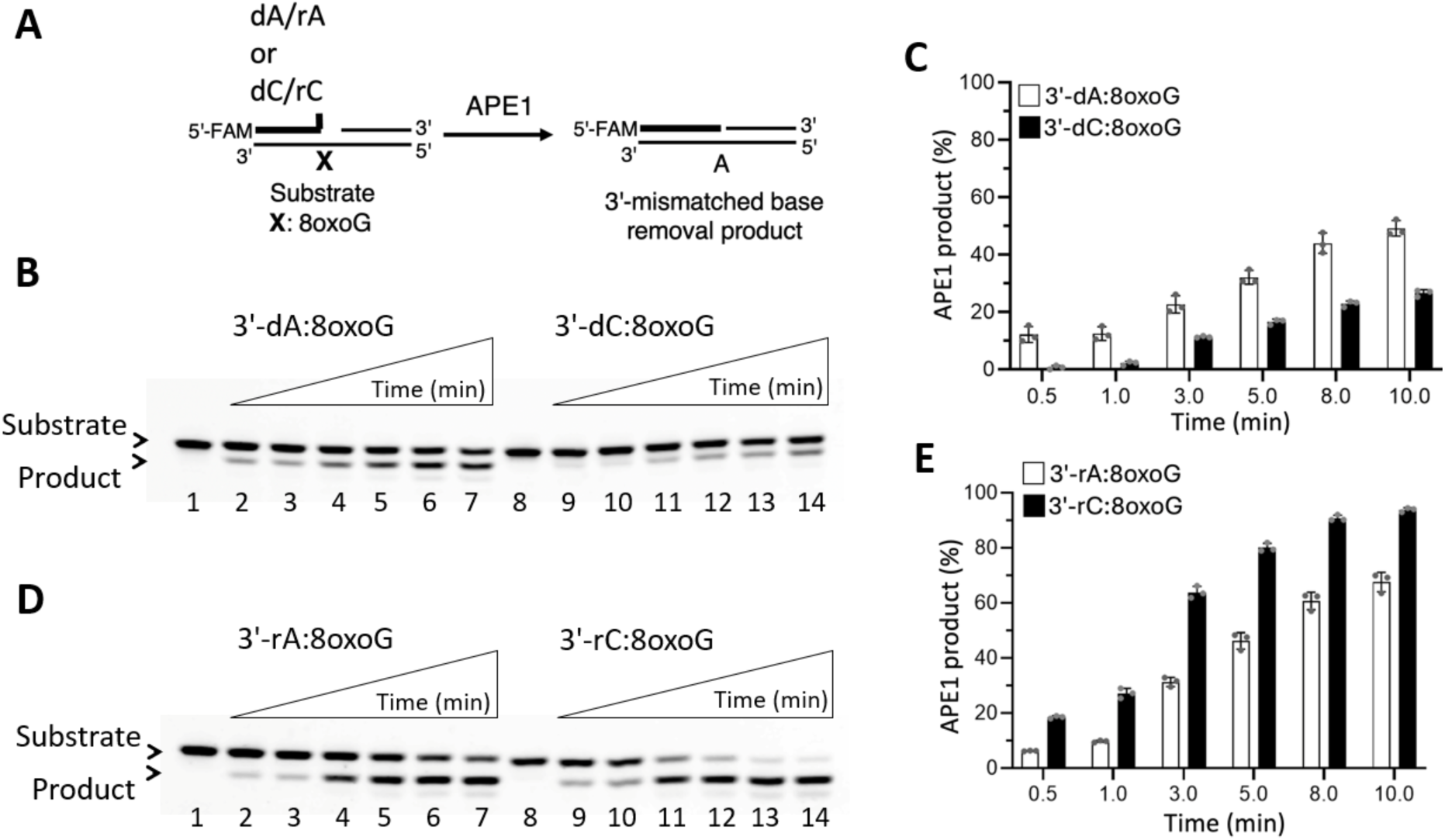
Mismatch removal by APE1 from the nick substrates with template 8-oxoG. **(A)** Scheme shows nick DNA substrates with a mismatch or a ribonucleotide and removal products by APE1 in exonuclease assay. (**B**) Lanes 1 and 8 are the negative enzyme controls of the nick DNA substrates with 3’-dA:8-oxoG and 3’-dC:8-oxoG, respectively. Lanes 2-7 and 9-14 are the removal of 3’-dA and 3’-dC mismatches from nick DNA substrates with 3’-dA:8- oxoG and 3’-dC:8-oxoG, respectively, and correspond to time points of 0.5, 1, 3, 5, 8, and 10 min. **(D)** Lanes 1 and 8 are the negative enzyme controls of the nick DNA substrates with 3’- rA:8-oxoG and 3’-rC:8-oxoG, respectively. Lanes 2-7 and 9-14 are the removal of 3’-rA and 3’-rC mismatches from nick DNA substrates with 3’-rA:8-oxoG and 3’-rC:8-oxoG, respectively, and correspond to time points of 0.5, 1, 3, 5, 8, and 10 min. **(C, E)** Graphs show time-dependent changes in the amount of mismatch (C) and ribonucleotide (E) removal products by APE1. The data represent the average of three independent experiments ± SD.

**Figure 16.**
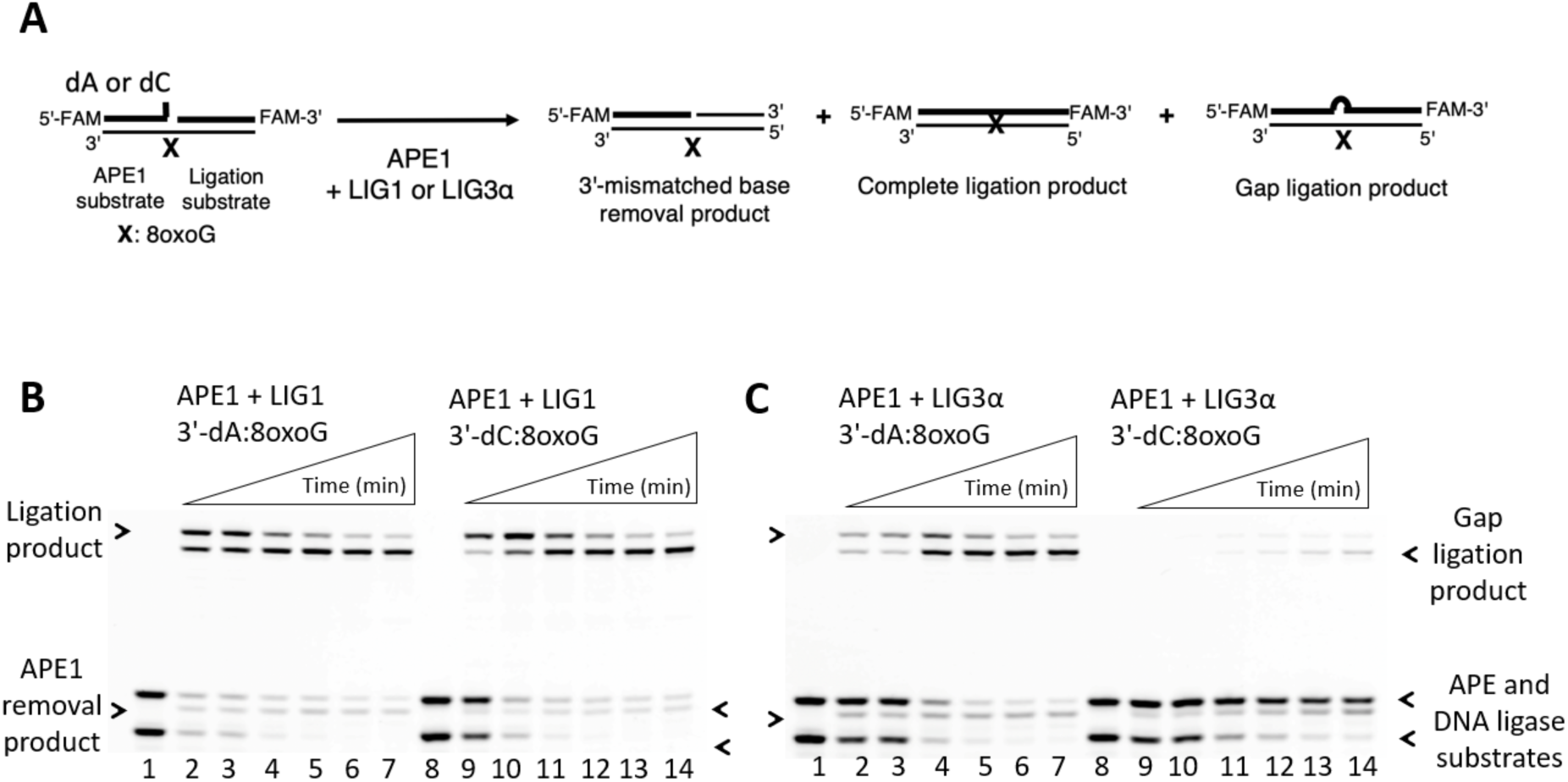
Ligation coupled to mismatch removal by APE1 and BER ligases in the presence of nick substrates with 3’-mismatches. **(A)** Scheme shows nick DNA substrates with 3’-mismatches and template 8-oxoG and products of ligation and mismatch removal by APE1 and BER ligases. **(B-C)** Lanes 1 and 8 are the negative enzyme controls of the nick DNA substrates with 3’-dA:8-oxoG and 3’-dC:8-oxoG, respectively. Lanes 2-7 and 9-14 are the coupled reaction products showing APE1 mismatch removal and ligation by LIG1 (B) and LIG3α (C) from the nick DNA substrates with 3’-dA:8-oxoG and 3’-dC:8-oxoG, respectively, and correspond to time points of 0.5, 1, 3, 5, 8, and 10 min.

Our results demonstrated that APE1 can remove 3’-dA and 3’-dC mismatches from nick substrates containing template 8-oxoG (Figure 15B-C), which is relatively less efficient than the removal of 3’-ribonucleotides (Figure 15D-E). In the presence of APE1 and the BER ligases (Figure 16A), the products of mismatch removal by APE1 and gap ligation after the removal are accumulated along with a complete ligation product over time in the presence of nick substrate with 3’-dA:8-oxoG (Figures 16B and 16C, lines 2-7). However, for nick substrate with 3’-dC:8-oxoG, we observed similar results only in the presence of LIG1 (Figure 16B, lanes 9-14) and no product was detected with LIG3α (Figure 16C, lanes 9-14). We also tested the interplay between APE1 and the BER ligases for nick substrates containing 3’- ribonucleotide and template 8-oxoG (Figure 17A). Overall, our results showed abundant gap ligation products in the presence of 3’-rA:8-oxoG and 3’-rC:8-oxoG, suggesting that both BER ligases can attempt to ligate gap repair intermediate after more efficient removal of 3’- ribonucleotides by APE1 (Figure 17B-C).

**Figure 17.**
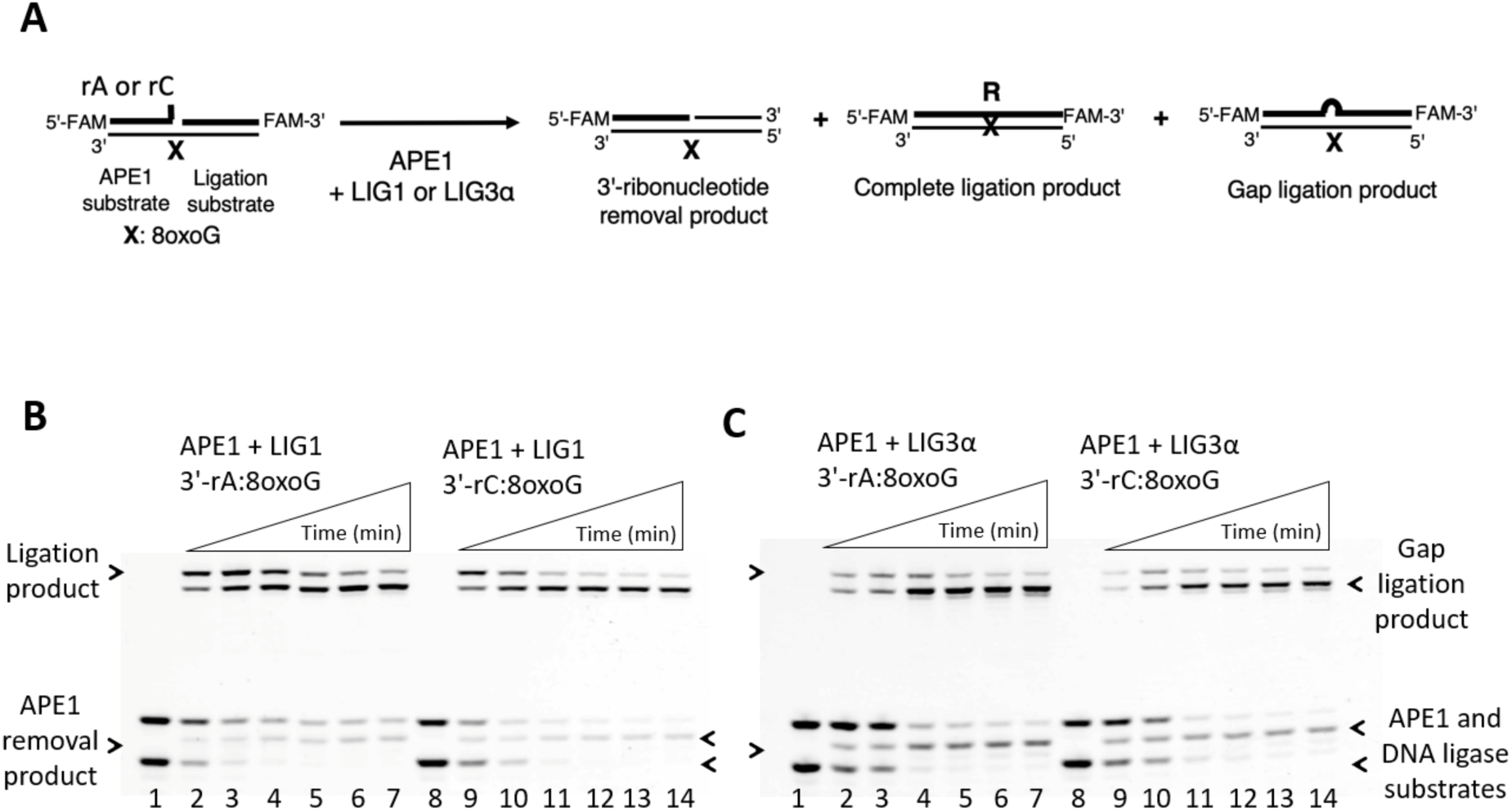
Ligation coupled to mismatch removal by APE1 and BER ligases in the presence of nick substrates with 3’-ribonucleotides. **(A)** Scheme shows nick DNA substrates with 3’-ribonucleotide and template 8-oxoG and products of ligation and a ribonucleotide removal by APE1 and BER ligases. **(B-C)** Lanes 1 and 8 are the negative enzyme controls of the nick DNA substrates with 3’-rA:8-oxoG and 3’-rC:8-oxoG, respectively. Lanes 2-7 and 9- 14 are the coupled reaction products showing APE1 ribonucleotide removal and ligation by LIG1 (B) and LIG3α (C) from the nick DNA substrates with 3’-rA:8-oxoG and 3’-rC:8-oxoG, respectively, and correspond to time points of 0.5, 1, 3, 5, 8, and 10 min.

## Discussion

Genomic DNA is constantly exposed to various endogenous sources and exogenous agents that generate the DNA base modifications (1). Among the most frequently formed DNA lesion created by oxidative stress, 8-oxoG not only arises spontaneously during normal cellular processes but is also generated by exposure to environmental sources such as UV radiation (2,3). The 8-oxoG lesions in the template strand could result in high frequency of misincorporation when pairs with either correct cytosine or incorrect adenine during replication and gap filling synthesis by DNA polymerases (4–6). Structural and kinetics studies have reported the reduced nucleotide selectivity of DNA polymerases at varying degrees from across families when they encounter template 8-oxoG relative to unmodified template guanine (7–10). This is mainly because of polymerase-specific variations in the ratio of dC:dA insertion opposite 8-oxoG and interactions at the polymerase active site in different conformations (13–16).

As the major cellular defense mechanism against oxidative DNA damage is BER (19), in the present study, we aimed to understand how the fidelity of 8-oxoG bypass by polβ in *anti*- vs *syn*- conformation could affect downstream steps involving final ligation of nick repair intermediate by LIG1 and LIG3α. In our previous studies, we characterized the molecular determinants of faithful BER and reported various types of deviations from the canonical repair pathway coordination during simultaneous activities of polβ and DNA ligases (23,32,33,41–46). Considering the insights gained from the current study, we propose a working model (Supplementary Scheme S1) that illustrates how polβ mismatch or ribonucleotide insertion during 8-oxoG bypass could impact the efficiency of subsequent nick sealing by LIG1 and LIG3α. We demonstrated that polβ inserting dATP in error-prone -*syn* conformation results in mutagenic ligation by both BER ligases leading to futile repair. However, while nick sealing efficiency of mutagenic dATP:8-oxoG insertion products was similar for both ligases, we observed a significantly diminished ligation by LIG3α after polβ dCTP:8oxoG insertion, demonstrating a substrate preference of BER ligase at the final step against polβ nick insertion products containing 8-oxoG lesion on template position in -*anti* vs -*syn* conformation. This could provide a fidelity check point for DNA-end processing enzymes, such as APE1, for proofreading of 3’-mismatched bases from nick repair intermediates.

The survey of the various ternary complex structures of polβ revealed that K280 side chain is flexible and can adopt multiple conformations (34–39). Our results demonstrated wild-type level of ligation efficiency by both BER ligases in case of mutagenic *syn*-conformation of template 8-oxoG in a Hoogsteen base pair with an incoming dATP by polβ K280A active site mutant. Furthermore, X-ray structures of polβ demonstrated that the Hoogsteen base pairing itself stabilizes the *syn*-conformation of 8-oxoG, but the enzyme also stabilizes the *syn*- conformation using hydrogen bonding between O8 of 8-oxoG and R283 side chain, along with an adjustment in the phosphodiester backbone of the template strand (36). Accordingly, we observed a complete ablation of mutagenic nick sealing after dATP:8-oxoG insertion by polβ R283K mutant, which is independent of BER ligase. This could be due to this increased fidelity opposite 8-oxoG by the effect of R283K mutation as shown in the ternary complex structure of the mutant with an incoming dATP (8). Polβ exhibits lower fidelity compared to other polymerases and lacks proofreading function (22). Moreover, 30% of human tumors have been found to express polβ variants carrying single amino acid substitutions resulting in aberrant function of the polymerase such as reduced fidelity or slow gap filling activity (47). We showed similar efficiencies of dATP:8-oxoG and dCTP:8-oxoG insertion by polβ E288K mutant and their subsequent ligation by both BER ligases. These findings suggest that the 8-oxoG- mediated mutagenesis during BER could be a contributor of the sequence specific mutator phenotype of polβ colon cancer-associated variant E288K as shown both *in vitro* and *in vivo* studies (48). Overall, these results provide an insight into the functional interplay between polβ and LIG1/LIG3α in case of alterations at the polymerase active site environment that affect the lesion bypass specificity of the polymerase leading to adjustments in the position of 3’-OH and 5’-PO4 ends of resulting nick repair intermediate that needs to be sealed by BER ligases at the final ligation step.

In addition to the ligation efficiency after polβ error-prone or -free insertions opposite 8-oxoG, we also showed the nick sealing ability of LIG1 and LIG3α favoring dA:8-oxoG mispair. Interestingly, the BER ligases exhibit a drastic difference for processing of 8-oxoG in -*anti* conformation that forms a Watson-Crick pair with cytosine. Our results showed inefficient ligation of polβ dCTP:8-oxoG insertion products and an inability to seal nick with 3’-dC:8- oxoG by LIG3α, demonstrating that the identity of BER ligase is an important determinant of faithful repair. Further structure studies are required to better elucidate the discrimination mechanism of LIG1 and LIG3α at atomic resolution against nick with templating 8-oxoG.

In addition to dNTP mismatches, we also demonstrated how ribonucleotide challenge could affect the ability of polβ bypassing the oxidative lesion and subsequent nick sealing step. Our results showed a discrimination against rATP and rCTP incorporation opposite 8-oxoG by polβ as the key factor in preventing the ligation of nick repair intermediates. We previously reported the role of this discrimination agaist rNTP insertions opposite undamaged bases for preventing ligation of repair intermediates (41). Overall results highlight a dramatic difference in the outcomes of the BER pathway coordination between polβ dNTP *versus* rNTP insertions coupled to ligation by LIG1 and LIG3α, specifically halting ligation when ribonucleotides are inserted across from template 8-oxoG and resulting in the ligation of one nucleotide gap repair intermediate. Despite the ribonucleotide’s ability to code for bases, a single nucleotide deletion could be more detrimental than an embedded one due to potential disruptions in the DNA sequence. However, it’s important to note that our results in the ligation assays with nick DNA substrates containing a single ribonucleotide and template 8-oxoG, *i.e*., preinserted 3’-rA:8- oxoG and 3’-rC:8-oxoG, showed an efficient nick sealing by LIG1 and LIG3α, demonstrating a lack of sugar discrimination. We also determined the impact LIG1 mutations that have been associated with immunodeficiency disease (49), found that the LIG1 syndrome variants similarly are more efficient in ligating ribonucleotide-containing nicks than those of nicks with 3’-mismatches in the presence of template 8-oxoG. Our findings showing a lack of sugar discrimination for ligating nicks are consistent with our previously solved structures of LIG1/RNA-DNA complexes, 3’-rG:C and 3’-rA:T (42). We reported that LIG1 can accommodate a single 3’-ribonucleotide at nick site mainly through a network of interactions between the ligase active sites D570 and R871, 2’-OH of the ribose, and water molecules leading to formation of final phosphodiester bond formation between DNA ends. Further structure studies are required to gain an atomic insight into how templating 8-oxoG in -*anti* vs -*syn* conformation at nick containing a 3’-ribonucleotide could affect sugar discrimination at the ligase active site. We previously reported the importance of physical interaction and functional interplay between APE1 and both ligases for BER efficiency (44). Accordingly, our results of the present study highlight this fidelity check point for BER accuracy at the downstream steps where APE1 removes 3’-mismatches or ribonucleotides from opposite template 8-oxoG of nick repair intermediates. We also showed that APE1 can coordinate with both ligases during proofreading coupled to nick sealing depending on the identity of 3’- mismatched base (dA/rA vs dC/rC) and BER ligase (LIG1 vs LIG3α).

In summary, the data presented here further contribute to elucidate the molecular determinants that dictate BER fidelity at the final steps where polβ coordinates with LIG1 and LIG3α during gap filling and subsequent nick sealing to finalize the repair process. We revealed that polβ mutagenic or error-free 8-oxoG lesion bypass, which could be formed by environmental oxidant-induced effects, could be an important source of mutagenic repair. To avoid toxicity, it is likely that these intermediates are quickly processed to fully repaired DNA and a block in the gap filling step and aberrant nick sealing subsequently could lead to the accumulation of stalled repair intermediates that could trigger the cell killing upon exposure to oxidative stress and genomic instability. Our findings demonstrated how the structural conformations that polβ shows depending on balance between the *anti*- and *syn*-conformations of 8-oxoG and the identity of an incoming nucleotide to be incorporated opposite the oxidative lesion could impact the repair efficiency at the downstream steps of BER pathway. Future studies in cells will be required to determine its precise impact to cytotoxicity to oxidative stress-inducing agents.

## Data Availability

Information and requests of materials used in this research should be directed to Melike Çaglayan.

## References

1. Evans, M.D., Dizdaroglu, M. and Cooke, M.S. (2004) Oxidative DNA damage and disease: Induction, repair and significance. Mutat. Res. Rev. Mutat. Res., 567, 1–61.

2. Nakabeppu, Y., Sakumi, K., Sakamoto, K., Tsuchimoto, D., Tsuzuki, T. and Nakatsu, Y. (2006) Mutagenesis and carcinogenesis caused by the oxidation of nucleic acids. Biol. Chem., 387, 373–379.

3. Marnett, L.J. (2000) Oxyradicals and DNA damage. Carcinogenesis, 21, 361–370.

4. Kouchakdjian, M., Bodepudi, V., Shibutani, S., Eisenberg, M., Johnson, F., Grollman, A.P. and Patel, D.J. (1991) NMR structural studies of the ionizing radiation adduct 7-hydro-8- oxodeoxyguanosine (8-oxo-7H-dG) opposite deoxyadenosine in a DNA duplex. 8-Oxo-7H- dG(syn).dA(anti) alignment at lesion site. Biochemistry, 30, 1403–1412.

5. Lipscomb, L.A., Peek, M.E., Morningstar, M.L., Verghis, S.M., Miller, E.M., Rich, A., Essigmann, J.M. and Williams, L.D. (1995) X-ray structure of a DNA decamer containing 7,8-dihydro-8-oxoguanine. Proc. Natl Acad. Sci. USA., 92, 719–723.

6. McAuley-Hecht, K.E., Leonard, G.A., Gibson, N.J., Thomson, J.B., Watson, W.P., Hunter, W.N. and Brown, T. (1994) Crystal structure of a DNA duplex containing 8- hydroxydeoxyguanine-adenine base pairs. Biochemistry, 33, 10266–10270.

7. Avkin, S. and Livneh, Z. (2002) Efficiency, specificity and DNA polymerase-dependence of translesion replication across the oxidative DNA lesion 8-oxoguanine in human cells. Mutat. Res./Fund. Mol. Mech. Mutagen., 510, 81–90.

8. Friedberg, E.C., Lehmann, A.R. and Fuchs, R.P. (2005) Trading places: how do DNA polymerases switch during translesion DNA synthesis? Mol. Cell.,18, 499–505.

9. Shibutani, S., Takeshita, M. and Grollman, A.P. (1991) Insertion of specific bases during DNA synthesis past the oxidation-damaged base 8-oxodG. Nature, 349, 431–434.

10. Brieba, L.G., Eichman, B.F., Kokoska, R.J., Doublie, S., Kunkel, T.A. and Ellenberger, T. (2004) Structural basis for the dual coding potential of 8-oxoguanosine by a high-fidelity DNA polymerase. EMBO J. 23, 3452–3461.

11. Al-Tassan, N. et al., (2002) Inherited variants of MYH associated with somatic G : C → T : A mutations in colorectal tumors. Nat. Genet., 30, 227–232.

12. Hsu, G.W., Ober, M., Carell, T. and Beese, L.S. (2004) Error-prone replication of oxidatively damaged DNA by a high-fidelity DNA polymerase. Nature, 431, 217–221.

13. Freisinger, E., Grollman, A.P., Miller, H. and Kisker, C. (2004) Lesion (in)tolerance reveals insights into DNA replication fidelity. EMBO J. 23, 1494–1505.

14. Shibutani, S., Takeshita, M. and Grollman, A.P. (1991) Insertion of specific bases during DNA synthesis past the oxidation-damaged base 8-oxodG. Nature, 349, 431–434.

15. Duarte, V., Muller, J.G. and Burrows, C.J.A.G. (1999) Insertion of dGMP and dAMP during in vitro DNA synthesis opposite an oxidized form of 7,8-dihydro-8-oxoguanine. Nucleic Acids Res., 27, 496–502.

16. McCulloch, S.D., Kokoska, R.J., Garg, P., Burgers, P.M. and Kunke, T.A. (2009) The efficiency and fidelity of 8-oxo-guanine bypass by DNA polymerases δ and η. Nucleic Acids Res., 37, 2840–2840.

17. Caldecott, K.W. (2020) Mammalian DNA base excision repair: Dancing in the moonlight. DNA Repair (Amst), 93, 102921.

18. Kim, Y.-J. and M. Wilson III, D. (2012) Overview of base excision repair biochemistry. Curr. Mol. Pharmacol., 5, 3–13.

19. Beard, W.A., Horton, J.K., Prasad, R. and Wilson, S.H. (2019) Eukaryotic base excision repair: New approaches shine light on mechanism. Annu. Rev. Biochem., 88, 137–162.

20. Wilson, S.H. and Kunkel, T.A. (2000) Passing the baton in base excision repair. Nature Structural Biology 7, 176–178.

21. Prasad, R., Shock, D.D., Beard, W.A. and Wilson, S.H. (2010) Substrate channeling in mammalian base excision repair pathways: passing the baton. J. Biol. Chem., 285, 40479.

22. Beard, W.A. and Wilson, S.H. (2006) Structure and mechanism of DNA polymerase β. Chem Rev 106, 361–382.

23. Çağlayan, M. (2019) Interplay between DNA Polymerases and DNA ligases: Influence on substrate channeling and the fidelity of DNA ligation. J. Mol. Biol., 431, 2068–2081.

24. Faucher, F., Doublié, S. and Jia, Z. (2012) 8-Oxoguanine DNA Glycosylases: One lesion, three subfamilies. Int. J. Mol. Sci., 13, 6711.

25. D’Augustin, O., Huet, S., Campalans, A. and Radicella, J.P. (2020) Lost in the crowd: How does human 8-Oxoguanine DNA Glycosylase 1 (OGG1) find 8-oxoguanine in the genome? 21, 8360.

26. Raetz, A.G. and David, S.S. (2019) When you’re strange: Unusual features of the MUTYH glycosylase and implications in cancer. DNA Repair (Amst), 80, 16–25.

27. Gordon, A.J.E., Satory, D., Wang, M., Halliday, J.A., Golding, I. and Herman, C. (2014) Removal of 8-oxo-GTP by MutT hydrolase is not a major contributor to transcriptional fidelity. Nucleic Acids Res., 42, 12015–12026.

28. Samaranayake, G.J., Huynh, M. and Rai, P. (2017) MTH1 as a Chemotherapeutic target: The elephant in the room. Cancers (Basel), 9, 47.

29. Tsuzuki, T., Nakatsu, Y. and Nakabeppu, K. (2007) Significance of error-avoiding mechanisms for oxidative DNA damage in carcinogenesis. Cancer Sci., 98, 465–470.

30. Nakabeppu, Y. (2001) Regulation of intracellular localization of human MTH1, OGG1, and MYH proteins for repair of oxidative DNA damage. Prog Nucleic Acid Res Mol Biol 68, 75–94.

31. Freudenthal, B.D., Beard, W.A., Perera, L., Shock, D.D., Kim, T., Schlick, T. and Wilson, S.H. (2015) Uncovering the polymerase-induced cytotoxicty of an oxidized nucleotide. Nature, 517, 635.

32. Çaǧlayan, M., Horton, J.K., Dai, D.P., Stefanick, D.F. and Wilson, S.H. (2017) Oxidized nucleotide insertion by pol β confounds ligation during base excision repair. Nat. Comm. 8, 1–11.

33. Çağlayan, M. and Wilson, S.H. (2015) Oxidant and environmental toxicant-induced effects compromise DNA ligation during base excision DNA repair. DNA Repair (Amst), 35, 85–89.

34. Freudenthal, B.D., Beard, W.A. and Wilson, S.H. (2013) DNA polymerase minor groove interactions modulate mutagenic bypass of a templating 8-oxoguanine lesion. Nucleic Acids Res., 41, 1848–1858.

35. Beard, W.A., Batra, V.K. and Wilson, S.H. (2010) DNA polymerase structure-based insight on the mutagenic properties of 8-oxoguanine. Mutat. Res., 703, 18–23.

36. Batra, V.K., Shock, D.D., Beard, W.A., McKenna, C.E. and Wilson, S.H. (2012) Binary complex crystal structure of DNA polymerase β reveals multiple conformations of the templating 8-oxoguanine lesion. Proc. Natl. Acad. Sci. U S A, 109, 113–118.

37. Batra, V.K., Beard, W.A., Hou, E.W., Pedersen, L.C., Prasad, R. and Wilson, S.H. (2010) Mutagenic conformation of 8-oxo-7,8-dihydro-2’-dGTP in the confines of a DNA polymerase active site. Nat. Struct. Mol. Biol., 17, 889–890.

38. Krahn, J.M., Beard, W.A., Miller, H., Grollman, A.P. and Wilson, S.H. (2003) Structure of DNA polymerase beta with the mutagenic DNA lesion 8-oxodeoxyguanine reveals structural insights into its coding potential. Structure, 11, 121–127.

39. Beard, W., Batra, V. and Wilson, S.H. (2010) DNA polymerase structure-based insight on the mutagenic properties of 8-oxoguanine. Mutat. Res., 703, 18–23.

40. Klein, H.L. (2017) Genome instabilities arising from ribonucleotides in DNA. DNA Repair (Amst), 56, 26–32.

41. Gulkis, M., Martinez, E., Almohdar, D. and Çaglayan, M. (2024) Unfilled gaps by polβ lead to aberrant ligation by LIG1 at the downstream steps of base excision repair pathway. Nucleic Acids Res., 52, 3810–3822.

42. Balu, K.E., Gulkis, M., Almohdar, D. and Çağlayan, M. (2024) Structures of LIG1 provide a mechanistic basis for understanding a lack of sugar discrimination against a ribonucleotide at the 3’-end of nick DNA. J. Biol. Chem., 300, 107216.

43. Balu, K.E., Tang, Q. Almohdar, D., Ratcliffe, J., Kalaycioglu, M. and Çağlayan, M. (2024) Structures of LIG1 uncover the mechanism of sugar discrimination against 5′-RNA-DNA junctions during ribonucleotide excision repair. J. Biol. Chem., 300, 107688.

44. Almohdar, D., Murcia, D., Tang, Q., Ortiz, A., Martinez, E., Parwal, T., Kamble, P. and Çağlayan, M. (2024) Impact of DNA ligase 1 and IIIα interactions with APE1 and polβ on the efficiency of base excision repair pathway at the downstream steps. J. Biol. Chem., 300, 107355.

45. Tang, Q., Gulkis, M., McKenna, R. and Çağlayan, M. (2022) Structures of LIG1 that engage with mutagenic mismatches inserted by polβ in base excision repair. Nat. Comm. 13, 1–11.

46. Çağlayan, M. (2020) The ligation of pol β mismatch insertion products governs the formation of promutagenic base excision DNA repair intermediates. Nucleic Acids Res., 48, 3708– 3721.

47. Sweasy, J.B., Lang, T., Starcevic, D., Sun, K.-W., Lai, C.-C., DiMaio, D. and Dalal, S. (2005) Expression of DNA polymerase cancer-associated variants in mouse cells results in cellular transformation. Proc. Natl. Acad. Sci. USA, 4, 14350–14355.

48. Murphy, D.L., Donigan, K.A., Jaeger, J. and Sweasy, J.B. (2012) The E288K colon tumor variant of DNA polymerase β is a sequence specific mutator. Biochemistry, 51, 5269– 5275.

49. Maffucci, P. et al. (2018) Biallelic mutations in DNA ligase 1 underlie a spectrum of immune deficiencies. J. Clin. Invest. 128, 5480–5504.

