## Supplementary Data for "Mutagenic ligation of polβ mismatch insertion products during 8-oxoG bypass by LIG1 and LIG3α at the downstream steps of base excision repair pathway"

### Supplementary Information

Supplementary Figures 1-14

Supplementary Tables 1-5

Supplementary Scheme 1

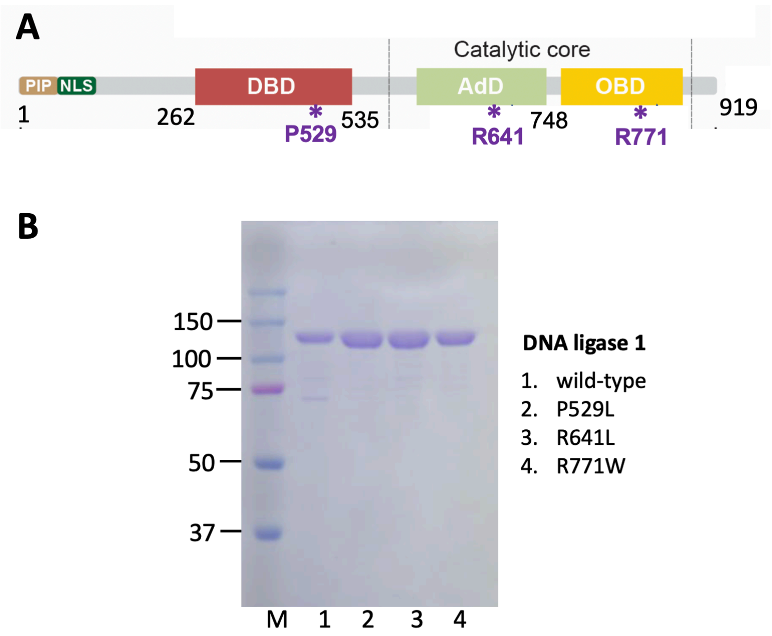


**Supplementary Figure 1**. **DNA ligase 1 proteins used in the study. (A)** Protein domain organization of human DNA ligase 1 (LIG1). N-terminal domain of LIG1 contains nuclear localization signal (NLS) and PCNA interacting box (PIP). C-terminal region of the protein includes catalytic core consisting of Oligonucleotide-Binding (OBD) and Adenylation (AdD) domains as well as DNA-binding (DBD) domain. The positions of the amino acid residues that are mutated in LIG1 deficiency disease are shown on the protein domain. **(B)** SDS-PAGE shows the purified LIG1 proteins wild-type and LIG1 deficiency disease-associated variants P529L, R641L, and R771W. M represents a Precision Plus Protein Dual Color Standard (10-250 kDa).

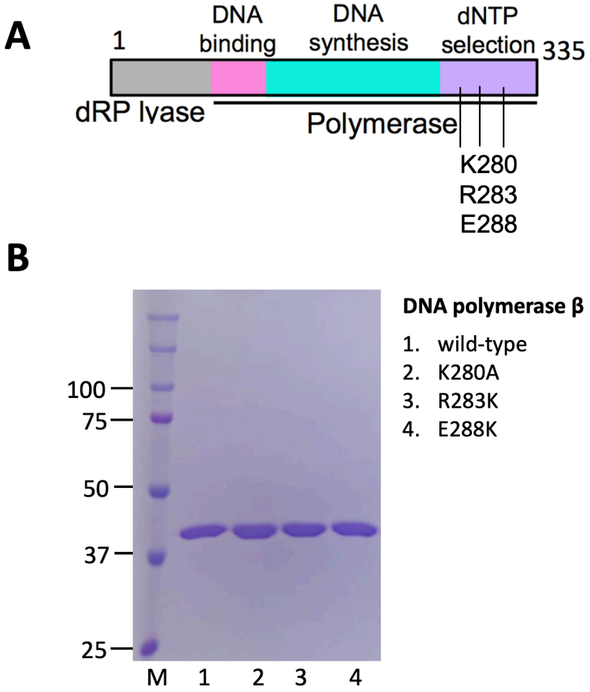


**Supplementary Figure 2**. **DNA polymerase β proteins used in the study. (A)** Protein domain organization of human DNA polymerase β (polβ). N-terminal part of the protein contains dRP-lyase domain that is responsible for the removal of 5'-dRP group and C-terminal region includes polymerase domain consisting of DNA binding, DNA synthesis, and dNTP selection sub-domains. The positions of the amino acid residues for polβ active site residues K280 and R283 as well as polβ cancer-associated mutation at E288 are shown on the protein domain. **(B)** SDS-PAGE shows the purified polβ proteins wild-type and mutants K280A, R283K, and E288K. M represents a Precision Plus Protein Dual Color Standard (10-250 kDa).

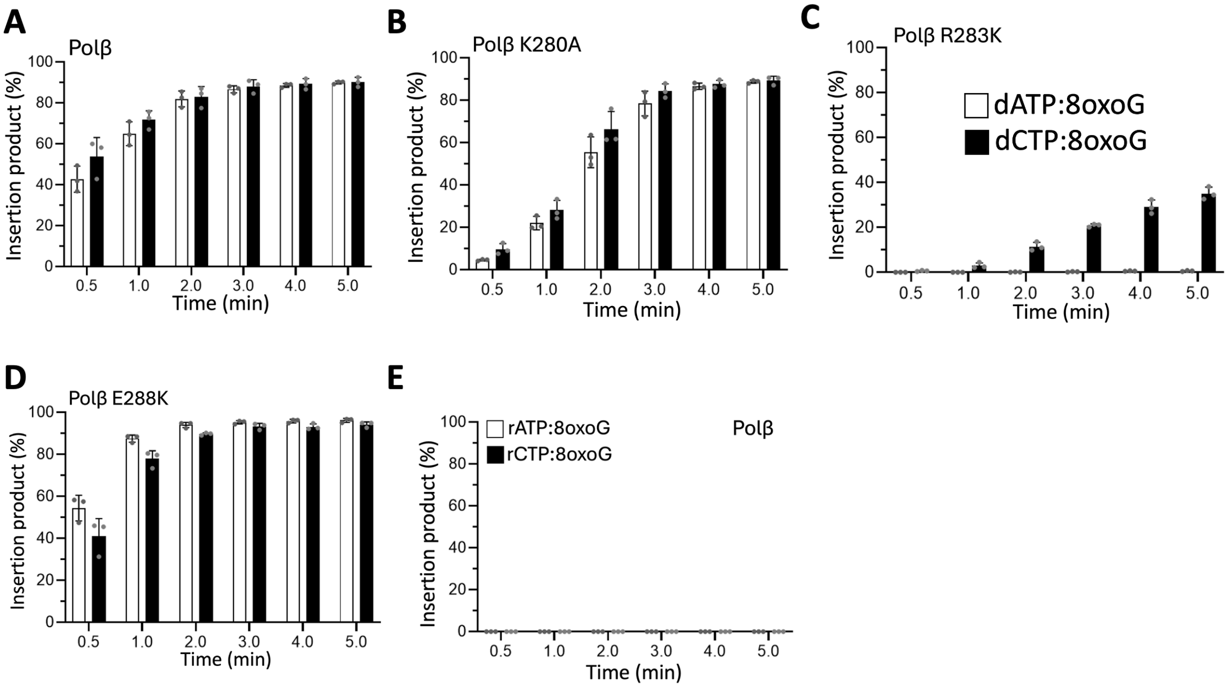


**Supplementary Figure 3. Comparison of mismatch nucleotide insertion products by polβ wild-type and mutants. (A-D)** Graphs show time-dependent changes in the amount of dATP and dCTP insertion products by polβ wild-type (A), active site mutants K280A (B), R283K (C), and cancer-associated variant E288K (D). **(E)** Graph shows time-dependent changes in the amount of rATP and rCTP insertion products by polβ wild-type. The data represent the average of three independent experiments ± SD.

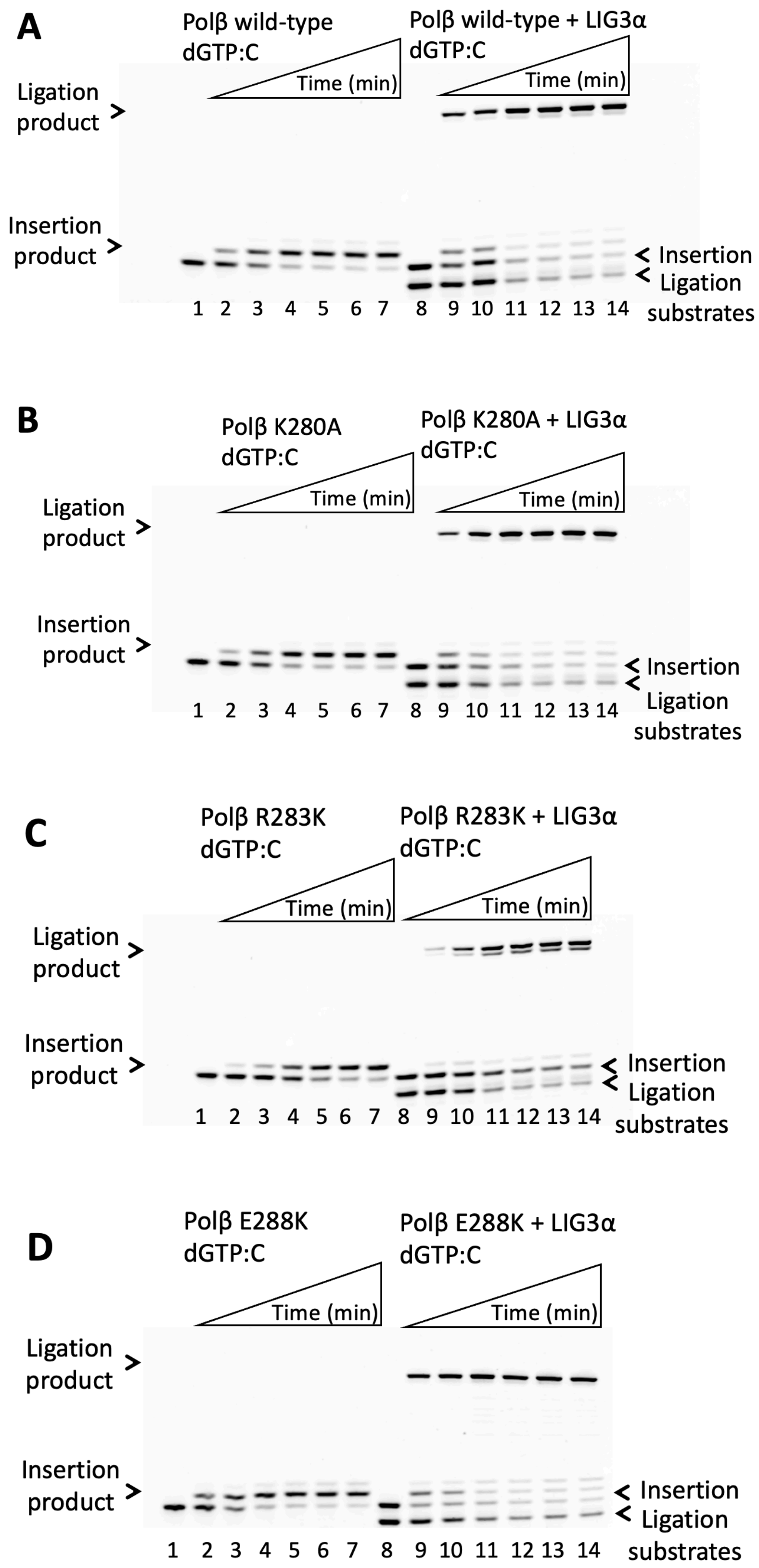
A
Polβ wild-type
dGTP:C
Time (min)
Polβ wild-type + LIG3α
dGTP:C
Time (min)
Ligation product
Insertion product
Insertion
Ligation
substrates
1 2 3 4 5 6 7 8 9 10 11 12 13 14
B
Polβ K280A
dGTP:C
Time (min)
Polβ K280A + LIG3α
dGTP:C
Time (min)
Ligation product
Insertion product
Insertion
Ligation
substrates
1 2 3 4 5 6 7 8 9 10 11 12 13 14
C
Polβ R283K
dGTP:C
Time (min)
Polβ R283K + LIG3α
dGTP:C
Time (min)
Ligation product
Insertion product
Insertion
Ligation
substrates
1 2 3 4 5 6 7 8 9 10 11 12 13 14
D
Polβ E288K
dGTP:C
Time (min)
Polβ E288K + LIG3α
dGTP:C
Time (min)
Ligation product
Insertion product
Insertion
Ligation
substrates
1 2 3 4 5 6 7 8 9 10 11 12 13 14

**Supplementary Figure 4. Ligation of correct dGTP:C insertion products by polβ wild-type and mutants. (A-D)** Lanes 1 and 8 are the negative enzyme controls of the one nucleotide gap DNA substrate with template C used in polβ insertion and polβ/DNA ligase coupled assays, respectively. Lanes 2-7 are dGTP:C insertion products by polβ wild-type (A), and mutants K280A (B), R283K (C), E288K (D), and correspond to time points of 0.5, 1, 2, 3, 4, and 5 min. Lanes 9-14 are the ligation of dGTP:C insertion products by polβ wild-type (A), K280A (B), R283K (C), E388K (D), and correspond to time points of 0.5, 1, 2, 3, 4, and 5 min.

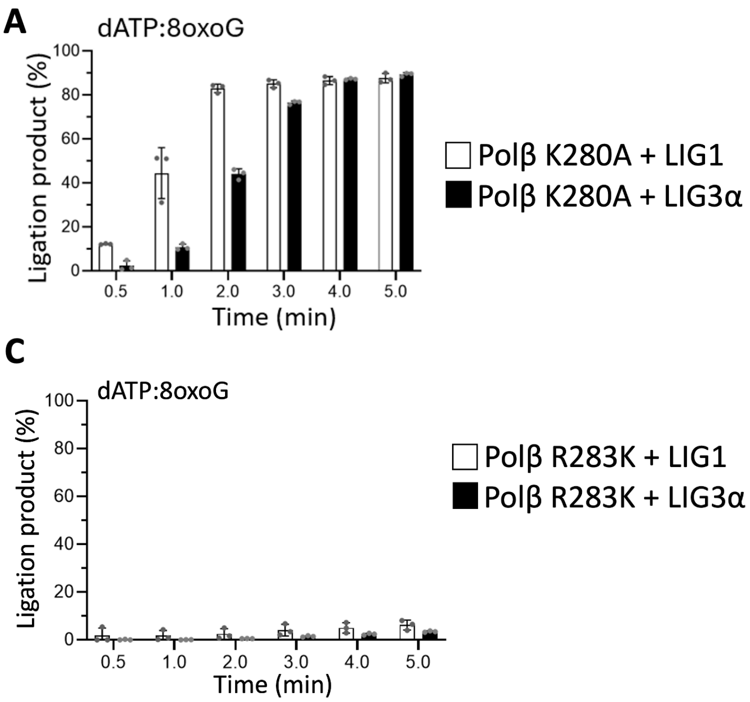


**Supplementary Figure 5. Comparison of ligation products by LIG1 *versus* LIG3α after dATP or dCTP insertion opposite template 8-oxoG by polβ active site mutants R280K and R283K. (A-B)** Graphs show time-dependent changes in the amount of ligation products after dATP:8oxoG (A) and dCTP:8oxoG (B) insertions by polβ K280A mutant to show the comparison between LIG1 and LIG3α. (**C-D)** Graphs show time-dependent changes in the amount of ligation products after dATP:8oxoG (C) and dCTP:8oxoG (D) insertions by polβ R283K mutant to show the comparison between LIG1 and LIG3α. The data represent the average of three independent experiments ± SD.

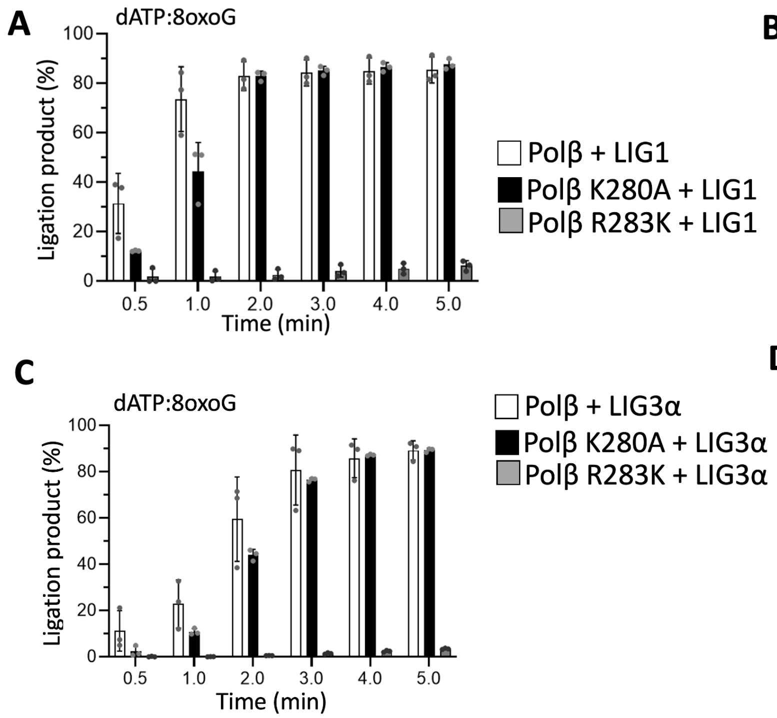


**Supplementary Figure 6. Comparison of ligation products by LIG1 or LIG3α after dATP or dCTP insertion opposite template 8-oxoG by polβ wild-type *versus* active site mutants R280K and R283K. (A-B)** Graphs show time-dependent changes in the amount of ligation products by LIG1 after dATP:8oxoG (A) and dCTP:8oxoG (B) insertions by polβ wild-type and active site mutants K280A and R283K. (**C-D)** Graphs show time-dependent changes in the amount of ligation products by LIG3α after dATP:8oxoG (C) and dCTP:8oxoG (D) insertions by polβ wild-type and active site mutants K280A and R283K. The data represent the average of three independent experiments ± SD.

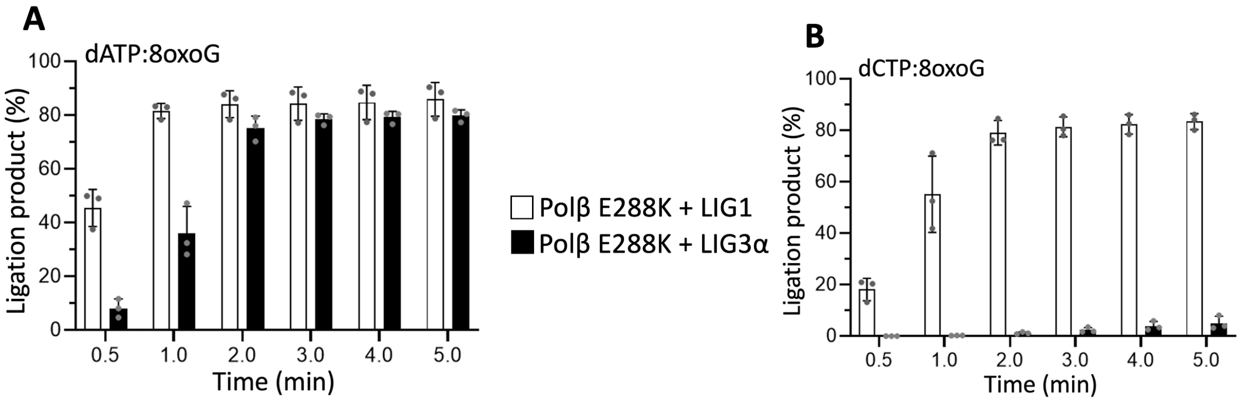


**Supplementary Figure 7. Comparison of ligation products by LIG1 *versus* LIG3α after dATP or dCTP insertion opposite template 8-oxoG by polβ cancer-associated variant E288K. (A-B)** Graphs show time-dependent changes in the amount of ligation products after dATP:8oxoG (A) and dCTP:8oxoG (B) insertions by polβ cancer-associated variant E288K to show the comparison between LIG1 and LIG3α. The data represent the average of three independent experiments ± SD.

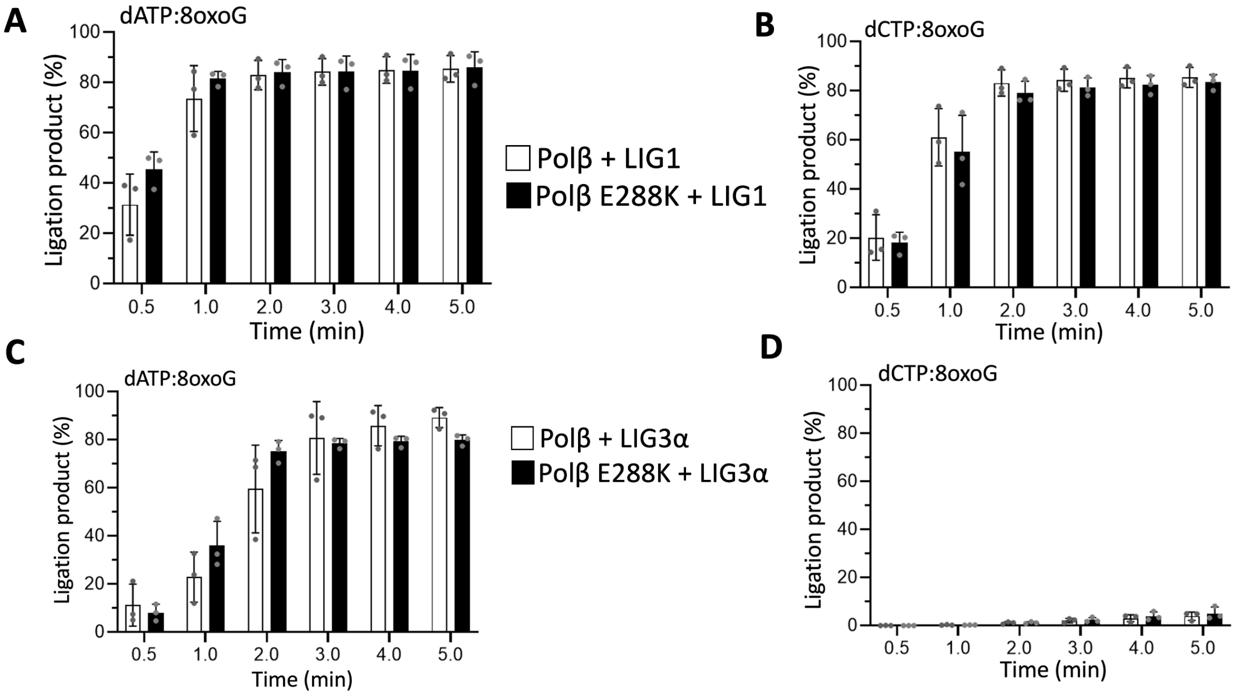


**Supplementary Figure 8. Comparison of ligation products by LIG1 or LIG3α after dATP or dCTP insertion opposite template 8-oxoG by polβ wild-type *versus* cancer-associated variant E288K. (A-B)** Graphs show time-dependent changes in the amount of ligation products by LIG1 after dATP:8oxoG (A) and dCTP:8oxoG (B) insertions by polβ wild-type and cancer-associated variant E288K. (**C-D)** Graphs show time-dependent changes in the amount of ligation products by LIG3α after dATP:8oxoG (C) and dCTP:8oxoG (D) insertions by polβ wild-type and cancer-associated variant E288K. The data represent the average of three independent experiments ± SD.

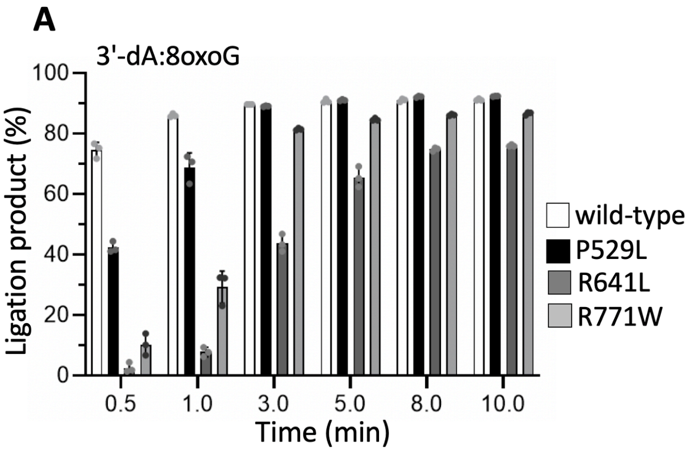


**Supplementary Figure 9. Comparison of ligation products by LIG1 wild-type and disease-associated variants. (A-B)** Graphs show time-dependent changes in the amount of ligation products by LIG1 wild-type *versus* LIG1 deficiency disease-associated variants P529L, R641L, R771W in the presence of nick DNA substrates with 3'-dA:8oxoG (A) and 3'-dC:8oxoG (B). The data represent the average of three independent experiments ± SD.

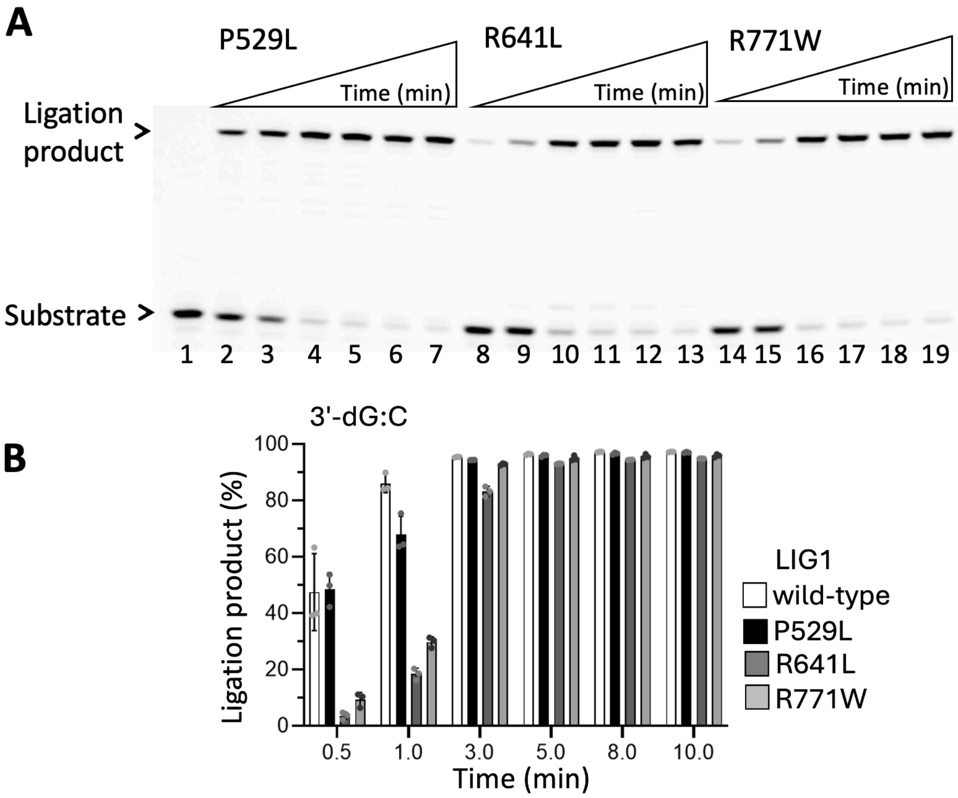


**Supplementary Figure 10. Ligation of nick DNA with canonical ends by LIG1. (A)** Line 1 is the negative enzyme control of the nick DNA substrate with 3'-dG:C. Lanes 2-7, 8-13, and 14-19 are the ligation products by LIG1 deficiency disease-associated variants P529L, R641L, and R771W, respectively, and correspond to time points of 0.5, 1, 3, 5, 8, and 10 min. **(B)** Graph shows time-dependent changes in the amount of ligation products by LIG1 wild-type *versus* LIG1 deficiency disease-associated variants P529L, R641L, R771W. The data represent the average of three independent experiments ± SD.

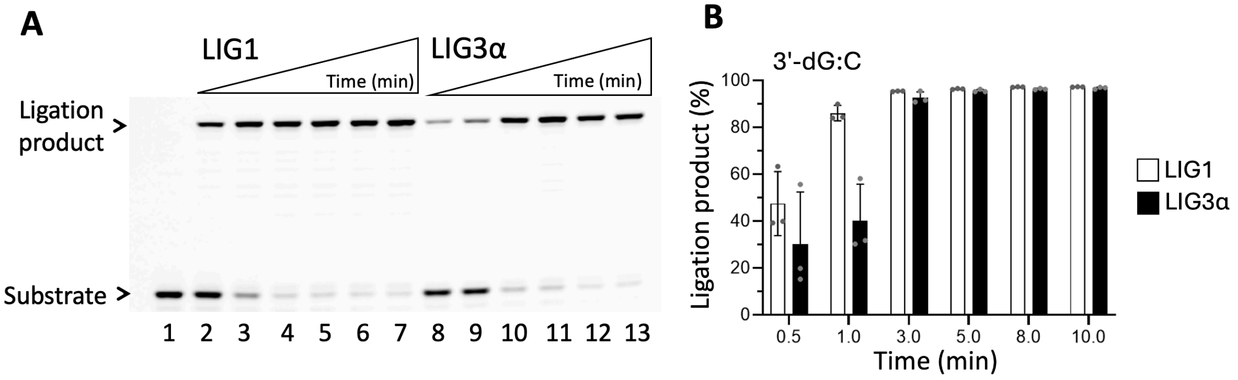


**Supplementary Figure 11. Ligation of nick DNA with canonical ends by LIG1 and LIG3α.** **(A)** Line 1 is the negative enzyme control of the nick DNA substrate with 3'-dG:C. Lanes 2-7 and 8-13 are the ligation products by LIG1 and LIG3α, respectively, and correspond to time points of 0.5, 1, 3, 5, 8, and 10 min. **(B)** Graph shows time-dependent changes in the amount of ligation products by LIG1 *versus* LIG3α, The data represent the average of three independent experiments ± SD.

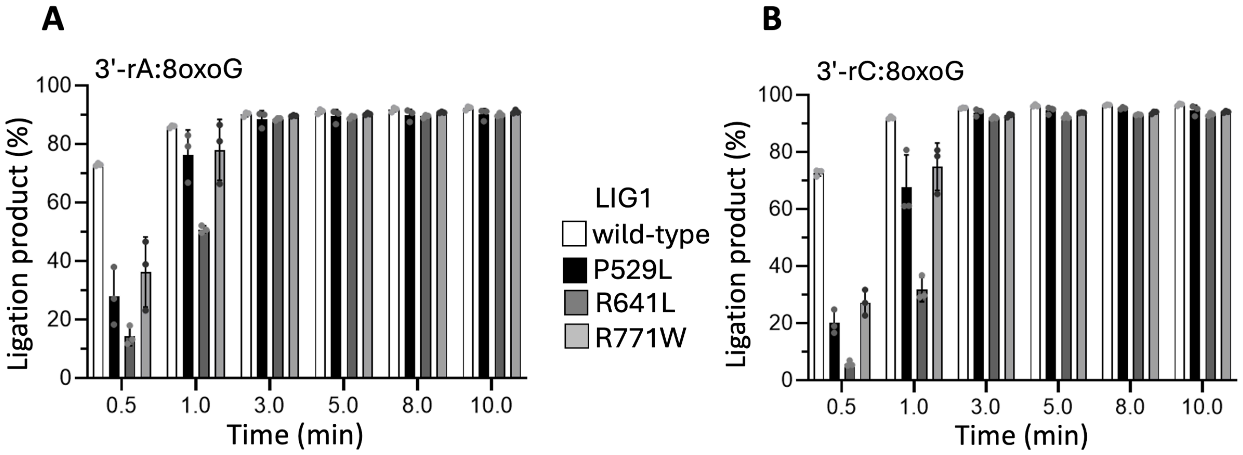


**Supplementary Figure 12. Comparison of ligation efficiency by LIG1 wild-type and deficiency disease-associated variants in the presence of nick DNA substrates with 3'-rA:8oxoG and 3'-rC:8oxoG. (A-B)** Graphs show time-dependent changes in the amount of ligation products by LIG1 wild-type *versus* LIG1 deficiency disease-associated variants P529L, R641L, R771W in the presence of nick DNA substrates with 3'-rA:8oxoG (A) and 3'-rC:8oxoG (B). The data represent the average of three independent experiments ± SD.

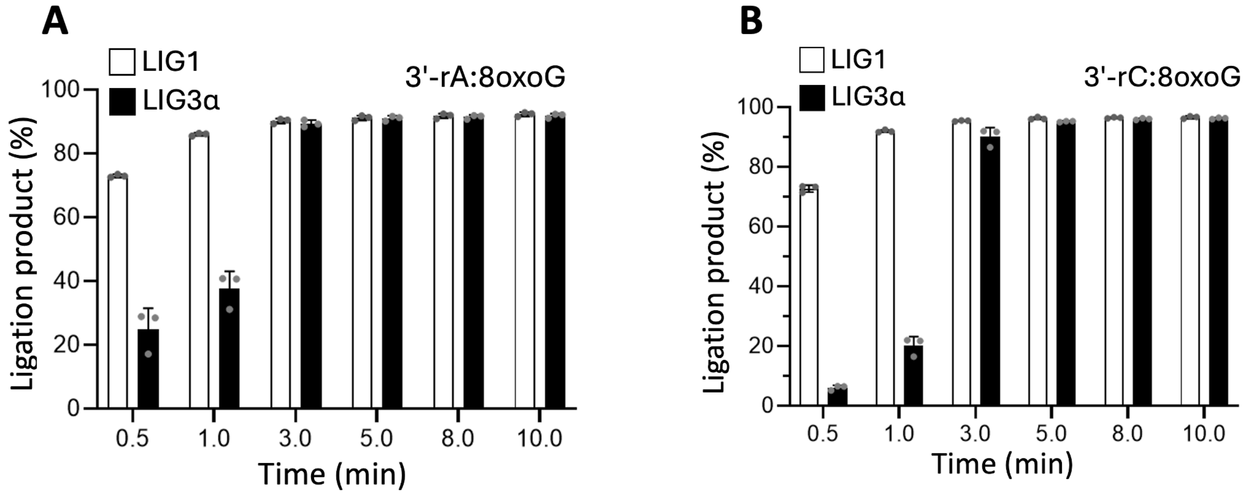


**Supplementary Figure 13. Comparison of ligation efficiency by LIG1 *versus* LIG3α in the presence of nick DNA substrates with 3'-rA:8oxoG and 3'-rC:8oxoG. (A-B)** Graphs show time-dependent changes in the amount of ligation products by LIG1 *versus* LIG3α in the presence of nick DNA substrates with 3'-rA:8oxoG (A) and 3'-rC:8oxoG (B). The data represent the average of three independent experiments ± SD.

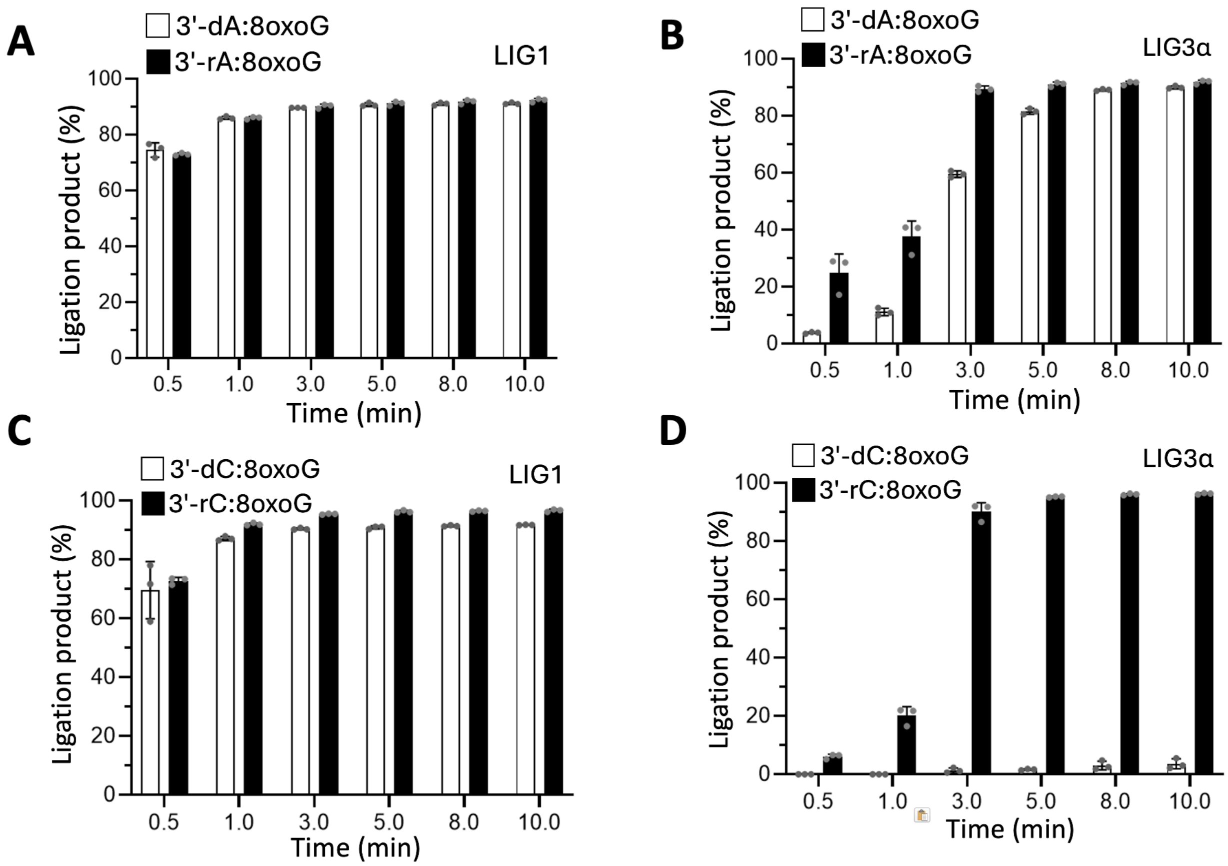


**Supplementary Figure 14. Comparison of ligation efficiency by LIG1 and LIG3α in the presence of nick DNA substrates with 3'-dA:8oxoG *versus* rA:8oxoG and 3'-dC:8oxoG *versus* rC:8oxoG. (A-D)** Graphs show time-dependent changes in the amount of ligation products and the data represent the average of three independent experiments ± SD.

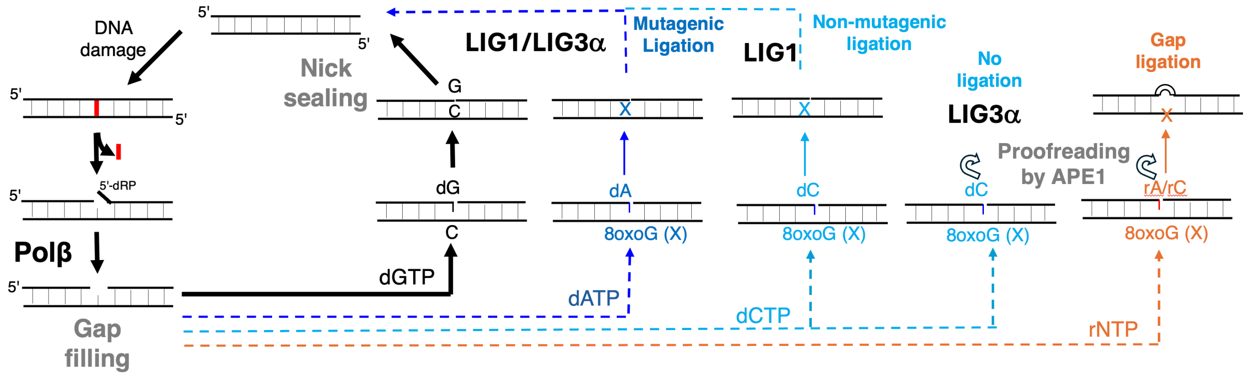


**Supplementary Scheme 1. Working model of the present study.** Illustration of the downstream steps of the BER pathway involving gap filling by polβ and subsequent nick sealing by LIG1 or LIG3α. In the presence of the correct nucleotide insertion by polβ (*i.e.*, dGTP:C), the nick repair product with proper ends (3'-OH and 5'-$PO_4$) can be efficiently ligated by both BER ligases. In the presence of 8-oxoG lesion on template position in -*syn* or -*anti* conformations, polβ bypassing the lesion by insertion of dATP mismatch opposite 8-oxoG leads to mutagenic ligation of resulting nick repair product by both LIG1 and LIG3α. Polβ dCTP insertion opposite template 8-oxoG results in non-mutagenic ligation by LIG1. Notably, the resulting nick repair product after dCTP:8oxoG insertion cannot sealed by LIG3α. However, in the presence of ribonucleotides, inability of polβ to insert rATP or rCTP opposite template 8-oxoG results in the ligation of gap repair intermediate by LIG1 and formation of single deletion mutagenesis products. APE1, via its proofreading role, can remove 3'-mismatches and 3'-ribonucleotides from nick repair intermediates containing 8-oxoG on a template position, thereby providing a fidelity check point at the final steps.

| DNA Substrates | Sequence |
| --- | --- |
| Gap DNA 8oxoG | FAM-5'-CATGGGCGGCATGAACC GAGGCCCATCCTCACC-3'<br>3'-GTACCCGCCGTACTTGGXCTCCGGGTAGGAGTGG-5' |
| Gap DNA C | FAM-5'-CATGGGCGGCATGAACC GAGGCCCATCCTCACC-3'<br>3'-GTACCCGCCGTACTTGGCCTCCGGGTAGGAGTGG-5' |

**Supplementary Table 1. Gap DNA substrates used in polβ nucleotide insertion assays.** One nucleotide gap DNA substrate with template base C or 8-oxoG and 5'-FAM label were used in the polβ insertion assays to test correct dGTP:C, mismatches dATP:8oxoG or dCTP:8oxoG, and ribonucleotides rATP:8oxoG or rCTP:8oxoG insertions. X presents 8-oxoG and FAM denotes a fluorescent tag. The base at the template position is underlined.

| DNA Substrates | Sequence |
| --- | --- |
| Gap DNA 8oxoG | FAM-5'-CATGGGCGGCATGAACC GAGGCCCATCCTCACC-3'-FAM<br>3'-GTACCCGCCGTACTTGGXCTCCGGGTAGGAGTGG-5' |
| Gap DNA C | FAM-5'-CATGGGCGGCATGAACC GAGGCCCATCCTCACC-3'-FAM<br>3'-GTACCCGCCGTACTTGGCCTCCGGGTAGGAGTGG-5' |

**Supplementary Table 2. Gap DNA substrates used in the coupled assays to measure gap filling by polβ and subsequent nick sealing by BER ligases.** One nucleotide gap DNA substrate with template base C or 8-oxoG and FAM labels at 3'- and 5'-ends were used in the coupled assays to test the ligation after polβ correct dGTP:C, mismatches dATP:8oxoG or dCTP:8oxoG, and ribonucleotides rATP:8oxoG or rCTP:8oxoG insertions by LIG1 or LIG3α. X presents 8oxoG and FAM denotes a fluorescent tag. The base at the template position is underlined.

| Nick DNA Substrates | Sequence |
| --- | --- |
| 3'-dG:C | 5'-CATGGGCGGCATGAACC**G**GAGGCCCATCCTCACC-3'-FAM<br>3'-GTACCCGCCGTACTTGGCCTCCGGGTAGGAGTGG-5' |
| 3'-dA:8oxoG | 5'-CATGGGCGGCATGAACC**A**GAGGCCCATCCTCACC-3'-FAM<br>3'-GTACCCGCCGTACTTGGXCTCCGGGTAGGAGTGG-5' |
| 3'-dC:8oxoG | 5'-CATGGGCGGCATGAACC**C**GAGGCCCATCCTCACC-3'-FAM<br>3'-GTACCCGCCGTACTTGGXCTCCGGGTAGGAGTGG-5' |
| 3'-rA:8oxoG | 5'-CATGGGCGGCATGAACC**rA**GAGGCCCATCCTCACC-3'-FAM<br>3'-GTACCCGCCGTACTTGGXCTCCGGGTAGGAGTGG-5' |
| 3'-rC:8oxoG | 5'-CATGGGCGGCATGAACC**rC**GAGGCCCATCCTCACC-3'-FAM<br>3'-GTACCCGCCGTACTTGGXCTCCGGGTAGGAGTGG-5' |

**Supplementary Table 3. Nick DNA substrates used in the ligation assays.** X presents 8-oxoG and FAM denotes a fluorescent tag. A ribonucleotide (rA or rC) or a mismatched base (dA or dC) at the 3'-end of nick DNA substrates are shown as bold and the template 8-oxoG is underlined.

| Nick DNA Substrates | Sequence |
| --- | --- |
| 3'-dA:8oxoG | FAM-5'-CATGGGCGGCATGAACC**A**GAGGCCCATCCTCACC-3'<br>3'-GTACCCGCCGTACTTGGXCTCCGGGTAGGAGTGG-5' |
| 3'-dC:8oxoG | FAM-5'-CATGGGCGGCATGAACC**C**GAGGCCCATCCTCACC-3'<br>3'-GTACCCGCCGTACTTGGXCTCCGGGTAGGAGTGG-5' |
| 3'-rA:8oxoG | FAM-5'-CATGGGCGGCATGAACC**rA**GAGGCCCATCCTCACC-3'<br>3'-GTACCCGCCGTACTTGGXCTCCGGGTAGGAGTGG-5' |
| 3'-rC:8oxoG | FAM-5'-CATGGGCGGCATGAACC**rC**GAGGCCCATCCTCACC-3'<br>3'-GTACCCGCCGTACTTGGXCTCCGGGTAGGAGTGG-5' |

**Supplementary Table 4. Nick DNA substrates used in APE1 exonuclease assays.** X presents 8-oxoG and FAM denotes a fluorescent tag. A ribonucleotide (rA or rC) or a mismatched base (dA or dC) at the 3'-end of nick DNA substrates are shown as bold and the template 8-oxoG is underlined.

| Nick DNA Substrates | Sequence |
|---|---|
| 3'-dA:8oxoG | FAM-5'-CATGGGCGGCATGAACC**A**GAGGCCCATCCTCACC-3'-FAM<br>3'-GTACCCGCCGTACTTGG$\underline{X}$CTCCGGGTAGGAGTGG-5' |
| 3'-dC:8oxoG | FAM-5'-CATGGGCGGCATGAACC**C**GAGGCCCATCCTCACC-3'-FAM<br>3'-GTACCCGCCGTACTTGG$\underline{X}$CTCCGGGTAGGAGTGG-5' |
| 3'-rA:8oxoG | FAM-5'-CATGGGCGGCATGAACC$^{r}$**A**GAGGCCCATCCTCACC-3'-FAM<br>3'-GTACCCGCCGTACTTGG$\underline{X}$CTCCGGGTAGGAGTGG-5' |
| 3'-rC:8oxoG | FAM-5'-CATGGGCGGCATGAACC$^{r}$**C**GAGGCCCATCCTCACC-3'-FAM<br>3'-GTACCCGCCGTACTTGG$\underline{X}$CTCCGGGTAGGAGTGG-5' |

**Supplementary Table 5. Nick DNA substrates used in the coupled assays to test APE1 exonuclease removal coupled to ligation by BER ligases.** X presents 8-oxoG and FAM denotes a fluorescent tag. A ribonucleotide (rA or rC) or a mismatched base (dA or dC) at the 3'-end of nick DNA substrates are shown as bold and the template 8-oxoG is underlined.
